# Phytic Acid (InsP_6_) Activates HDAC3 Epigenetic Axis to Maintain Intestinal Barrier Function

**DOI:** 10.1101/2024.09.15.613154

**Authors:** Sujan Chetterjee, Zachary Sin, Nguyen Tran, Loretta Vierra, Anuj Shukla, Tam Tran, George Koshkaryan, Kevin Ritter, Xue Bessie Su, Saharat Jolak Ragsac, Seungman Park, Qian Liu, Richard Van, Mira Han, Adolfo Saiardi, Katherine Huang, Kayci Huff-Hardy, Richard Rood, Anas Gremida, Martin Gregory, Chien-Huan Chen, Parakkal Deepak, Henning J. Jessen, Prasun Guha

## Abstract

While HDAC inhibition shows promise in cancer treatment, pan-HDAC inhibitors cause gastrointestinal issues in 48% of patients. Understanding HDAC activation mechanisms is crucial to treat diverse diseases beyond cancer. Our study reveals the essential role of inositol polyphosphate multikinase (IPMK) and inositol hexakisphosphate (InsP_6_ or phytic acid), enriched in vegan diets, in activating the HDAC3 epigenetic axis and maintaining intestinal barrier integrity. IPMK binds to HDAC3, driving InsP_6_ synthesis, which selectively activates HDAC3 at 10nM concentration by recruiting the DAD domain of its corepressor protein. *IPMK* deletion diminishes HDAC3 activation, leading to histone hyperacetylation and MMP gene transcription, compromising intestinal barrier integrity. InsP_6_ treatment is sufficient to rescue these effects. In inflammatory bowel disease, diminished IPMK levels exacerbated intestinal permeability, while oral InsP_6_ treatment mitigated gut permeability by restoring the HDAC3 epigenetic axis, indicating the clinical implications of the IPMK-HDAC3 epigenetic axis and therapeutic potential of phytic acid.

## Introduction

HDAC3, a key regulator of histone acetylation^1^, plays a crucial role in epigenetic modification^1^. While nonspecific HDAC inhibitors show promise in cancer treatment ^2^, their side effects^3^ highlight the importance of understanding individual HDAC functions. HDAC3, a prominent class I HDAC family member^1^, has emerged as a critical regulator of various physiological functions. Preclinical mouse model studies have demonstrated that HDAC3 is pivotal in maintaining metabolic homeostasis and circadian rhythms, particularly in the liver^4^. Furthermore, HDAC3 loss has been observed in inflammatory bowel disease (IBD) patients^5^, and its deletion in mouse intestinal epithelial cells induces Crohn’s disease-like symptoms, including increased intestinal permeability^5^. Despite its physiological importance, our understanding of the underlying mechanisms regulating HDAC3’s enzymatic activity remains limited.

What is known about the HDAC3 activation mechanism is mostly related to protein-protein interaction, suggesting HDAC3 binds to the corepressor complex silencing mediator for retinoid and thyroid hormone receptors (SMRT/NCoR2)^1^ and nuclear receptor co-repressor 1 (NCoR1)^1^, essential for its enzymatic activity^1^. Recent studies have illuminated the interaction between HDAC3 and inositol phosphates, particularly InsP_4,_ through crystal structure analysis^1^. However, significant gaps in our understanding persist. The physiological basis of InsP’s role in activating HDAC3 remains unclear. The precise mechanism and cellular location (cytoplasm, nucleoplasm, or chromatin) of InsP-mediated HDAC3 activation are unknown. Most crucially, the downstream effects of InsP-activated HDAC3 on the epigenetic axis, gene regulation, and overall cell function remain largely unclear.

While negatively charged InsP_4_ binding to HDAC3 was charge-dependent^1,6^, in a cellular environment, it is produced in very small amounts^7^. In contrast, InsP_6_, which is more negatively charged than InsP_4_, is produced in cells at higher micromolar concentrations and is more abundant^8^. Although the crystal structure did not observe InsP_6_ binding to HDAC3^1^, biochemical *in vitro* studies identified high-affinity binding of InsP_6_ to HDAC3^6^. While *in vitro* studies show InsP_4_, InsP_5_, and InsP_6_ can activate purified HDAC3 complexes from wild-type cells, these experiments involved dissociating InsPs in high-salt conditions^6,9^. Although this suggests InsPs regulate HDAC3 outside the cellular environment, it doesn’t confirm that physiological InsPs influences HDAC3 function. Thus, the biological significance of *in vivo* InsP-mediated HDAC3 regulation remains unclear.

At the physiological level, intracellular InsPs production is regulated by several enzymes, with IPMK (inositol polyphosphate multikinase) playing a critical role^8^, as its deletion diminishes InsP_4_, InsP_5_, and InsP_6_ levels^7^. Here, we investigated the effect of IPMK loss of function on HDAC3 activation to determine the importance of InsPs in regulating HDAC3 activation and the precise molecular mechanism. We discovered that IPMK physically interacts with HDAC3, and the interaction occurs specifically in chromatin but not in other cellular compartments. This finding is particularly intriguing given the known dynamic behavior of HDAC3, which shuttles between the cytoplasm and nucleus and requires recruitment to chromatin for its deacetylation function^10^. This chromatin-specific binding of IPMK to HDAC3 suggests a targeted regulatory mechanism, potentially influencing HDAC3’s activity precisely where it exerts its epigenetic effects.

Our research further uncovered a critical role for InsP_6_ in the activation of HDAC3. Notably, treatment with InsP_6_ to *IPMK* knockout cells and mouse models completely restored HDAC3 activity to levels comparable to those observed in wild-type cells, firmly establishing InsP_6_ as a physiological small molecule activator of HDAC3.

Notably, InsP_6_, also known as phytic acid, is abundant in plant-based diets, particularly in rice and legumes^11^. This dietary connection becomes significant in light of our findings. We discovered that in a mouse model of inflammatory bowel disease (IBD), there was a notable loss of IPMK protein expression. This loss disrupted the IPMK-HDAC3 epigenetic axis, leading to the acetylated histone lysine 16 (H4K16) promoter enrichment and transcriptional upregulation of matrix metalloproteinase (MMP) genes elevating gut permeability. MMPs are proteolytic enzymes that play a crucial role in degrading cell-cell junctions, consequently exacerbating intestinal permeability, a hallmark feature of IBD^12^. This aligns with genome-wide association studies (GWAS) and our preclinical study that have identified IPMK as a risk gene for IBD^13^. While transcriptional upregulation of MMPs in IBD is well-established^12,14^, the mechanism of epigenetic regulation of their expression was previously obscure. Our findings suggest that disruption of the IPMK-HDAC3 epigenetic axis influence the transcriptional upregulation of MMP family genes in IBD. Notably, we demonstrate that oral InsP_6_ (Phytic acid) treatment is effective in reducing intestinal permeability. .

In conclusion, our study reveals that IPMK physically interacts with chromatin-bound HDAC3 and identifies InsP_6_ as a crucial physiological small molecule activator of HDAC3. InsP_6_ functions by recruiting the DAD domain of the corepressor to HDAC3, thereby activating it and leading to the deacetylation of specific histone marks. This mechanism plays a crucial role in modulating the epigenetic landscape, specifically by repressing MMPs transcription. Consequently, it helps maintain cell-cell junctions and preserve intestinal barrier integrity, highlighting its importance in gut homeostasis and potentially in preventing or intercepting inflammatory bowel diseases.

## Results

### IPMK binds to the nuclear HDAC3 and is essential for HDAC3 activation regulating histone acetylation

To investigate the physiological impact of HOIPs on HDAC3 activation, *IPMK* was deleted in HCT116 (colon), A549 (lung), and MEF (mouse embryonic fibroblast cell lines **(Extended Fig1 A)**. As the rate-limiting enzyme in the HOIP pathway **(Fig 1A)**, *IPMK* deletion is expected to cause a profound loss in the levels of all downstream HOIP pathway products^8^. Consistent with this expectation, we observed a significant reduction in the levels of InsP_5_ and InsP_6_ in *IPMK* knockout (KO) cells, with only a modest decline in InsP_4_ **(Fig 1B)**. Additionally, HPLC data from the wild-type (WT) cells indicated that the overall concentrations of InsP_5_ and InsP_6_ are 3 to 4 times higher than that of InsP_4_ **(Fig 1B)**. To investigate the impact of *IPMK* deletion on HDAC3’s enzymatic activity, we immunopurified endogenous HDAC3 from respective WT and *IPMK* KO cells, followed by HDAC3 activity assay **(Fig 1C)**. We optimized the HDAC3 activity assay by immunoprecipitating HDAC3 from various protein concentrations ranging from 5μg, 20μg, 100μg and 500μg. Our results showed that HDAC3 purified from 20μg of protein exhibited strong deacetylase activity, with only modest increments in activity observed up to 500μg **(Extended Fig 1B)**. Based on these findings, we selected 20μg as the minimum protein concentration for all subsequent HDAC3 pulldown followed by enzymatic assays. This choice allowed us to achieve better resolution in detecting activity differences. Notably, *IPMK* deletion dramatically reduced HDAC3’s enzymatic activity by over 90% in all tested cell lines compared to their WT counterparts **(Fig1 D),** however, without affecting HDAC3 or corepressor protein levels **(Extended Fig 1C)**. Strikingly, this reduction in HDAC3 activity was comparable to that observed in *HDAC3*-KO cells **(Extended Fig 1D)** or WT cells treated with the HDAC3-specific inhibitor RGFP966 **(Fig1 E).** The selectivity of RGFP966 for HDAC3 is precise, as it shows 200-fold selectivity for HDAC3 compared to other HDACs and binds to HDAC3’s enzymatic site, inhibiting its deacetylase activity^15^. These findings demonstrate that at the physiological level, *IPMK* deletion impairs HDAC3 activation to an extent similar to complete *HDAC3* deletion or pharmacological inhibition, highlighting IPMK’s crucial role in maintaining HDAC3 function.

**Figure 1:**
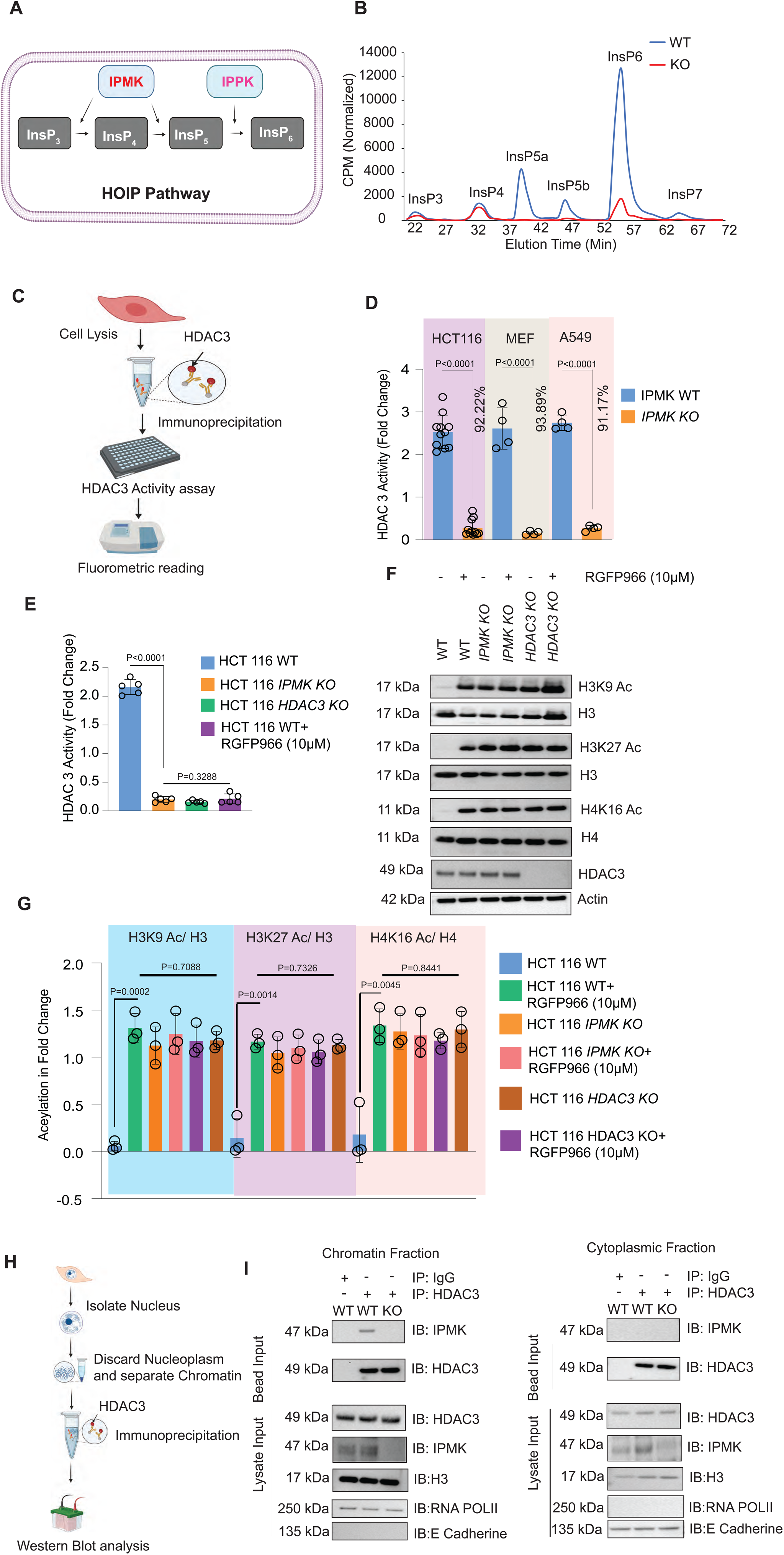
IPMK binds chromatin-bound HDAC3, regulating its activation and associated epigenetic changes. **(A)** Pictorial representation demonstrates the function of IPMK in the higher-order inositol phosphate (HOIP) pathway. IPMK converts InsP_3_ to InsP_4_ and InsP_4_ to InsP_5_, while IPPK converts InsP_5_ to InsP_6_. **(B)** HPLC analysis of inositol phosphate level depicted *IPMK* deletion reduced InsP_5_ and InsP_6_ levels in HCT 116 cells. The blue line indicated Wild type HCT116 cells (WT), and the red line indicated *IPMK-* Knock out HCT 116 cells (KO). Result was representative of two individual experiments (n=2). **(C)** Schematic diagram of HDAC3 activity assay. Immunoprecipitation of HDAC3 from cell lysate followed incubation with substrate and measuring HDAC3 activity fluorometrically. **(D)** HDAC3 were immunoprecipitated from Wild Type (WT) and *IPMK-* Knock out (KO) HCT116, MEF and A549 cells followed by HDAC3 activity assay. HDAC3 activity was markedly decreased in *IPMK-*Knock out (KO) HCT116, MEF and A549 cell lines compared to respective Type (WT) cells (HCT 116 n=12, MEF n=4, A549 n=4). Data was represented as a fold change compared to IgG. Results are representative of minimum four individual experiments. Percentage decrease of HDAC3 activity in IPMK KO cells compared to the WT was shown respective to each cell line. **(E)** Deletion of *IPMK* diminished HDAC3 activity in HCT 116 cells which is comparable to CRISPR-knock out *HDAC3* cells or WT cells treated with the HDAC3-specific inhibitor RGFP966 (10 µM for 24 hrs). Data was represented as a fold change compared to IgG. Results are representative of minimum three individual experiments (n=3). **(F)** Western blot analysis revealed that the deletion of *IPMK* or *HDAC3* led to profound increased acetylation of H3K9, H3K27, and H4K16 compared to wild-type cells. The levels of acetylation observed in both mutant cell types were comparable. An 8-hour treatment with the HDAC3-specific inhibitor RGFP966 at a dose of 10 µM in HCT116 WT, *IPMK* KO, and *HDAC3* KO cells showed no further increase in H3K9, H3K27, and H4K16 acetylation compared to the untreated *IPMK* KO and *HDAC3* KO cells. Lysates were immunoblotted against acetylated anti-H3K9, anti-H3k27, anti-H4k16 and HDAC3 antibodies and followed by stripping and reprobing against anti-H3 or anti-H4 antibody and actin, respectively, used as loading controls. Data is representative of three experimental replicates (n=3). **(G)** Representative densitometric analysis of H3k9ac / H3, H3k27ac / H3 and H4k16ac / H3. (n=3). **(H)** Schematic diagram of HDAC3 chromatin immuno precipitation. **(I)** HDAC3 was immunopurified using an anti-HDAC3 antibody from the chromatin fraction, and cytoplasmic fraction, followed by western blot analysis with an anti-IPMK antibody confirmed that IPMK exclusively binds to HDAC3 in chromatin fraction but not with cytoplasmic HDAC3. IgG was used as a negative control. Immunoblotting against anti-HDAC3, anti-IPMK, anti-H3 antibodies from the chromatin fraction isolated from *IPMK-* Wild type HCT116 (WT) and *IPMK-* Knock out HCT 116 (KO) cells were used as input control while anti-RNA pol II and anti-E-Cadherin used as cell fractionation control. Immunoblot with anti-HDAC3 from the bead used as loading control. The result was representative of three individual experiments (n=3).

Given HDAC3’s role as a deacetylase, the loss of its activity in *IPMK* KO cells was expected to increase histone acetylation. Our findings confirmed that *IPMK* deletion enhanced acetylation levels of specific histone residues, particularly H3K9/27 and H4K16 **(Extended Fig 1E, F)**. Notably, the extent of H3K9/27 and H4K16 acetylation in *IPMK* KO cells was comparable to that observed in *HDAC3* KO cells and wild-type cells treated with the HDAC3 inhibitor RGFP966 **(Fig 1 F, G)**. To determine whether this effect in *IPMK* deleted cells was due to the loss of HDAC3 activity, we treated *IPMK* KO cells with the HDAC3-specific inhibitor RGFP966. Interestingly, the inhibitor failed to induce any further increase in histone acetylation either in *IPMK* or *HDAC3* KO cells **(Fig 1 F, G)** while profoundly increasing H3K9/27 and H4K16 acetylation when treated to WT cells **(Fig 1 F, G)**. This differential response strongly suggests that HDAC3 activity was already substantially compromised in *IPMK*-deficient cells; consequently, RGFP966 was unable to enhance histone acetylation in *IPMK* KO cells further.

Next, we explored the potential direct interaction between IPMK and HDAC3 under physiological conditions. Considering HDAC3’s known shuttling between the nucleus and cytoplasm^10^ and its requirement to bind chromatin for histone deacetylation in the nucleus, we performed cell fractionation and chromatin immunopurification, revealing that IPMK specifically associates with chromatin-bound HDAC3 **(Fig 1 H, I)**. We did not find the interaction between IPMK and HDAC3 in the cytoplasm; however, it was detected when chromatin-bound HDAC3 was immunopurified **(Fig 1H, I)**. To further validate this interaction, we conducted an *in vitro* binding assay using recombinant HDAC3 and IPMK proteins, which confirmed a direct physical interaction between these two proteins **(Extended Fig 1G)**.

Overall, the study demonstrated that IPMK binds to nuclear HDAC3 and is crucial for its enzymatic activity.

### IPMK’s inositol kinase activity is essential for HDAC3 activation

To elucidate the role of IPMK’s inositol kinase activity in regulating HDAC3’s enzymatic activity, we conducted experiments using *IPMK* knockout (KO) cells. We transfected these cells with either wild-type (WT) or kinase-dead (KD) IPMK^7^. Our results showed that rescuing *IPMK* KO cells with WT-IPMK restored HDAC3 deacetylase activity, while KD-IPMK failed to do so **(Fig 2A)**. Interestingly, immunoprecipitation studies revealed that both WT-IPMK and KD-IPMK could interact with HDAC3 complex proteins **(Fig 2B)**. These findings suggest a crucial distinction: while IPMK’s physical binding to HDAC3 is insufficient for rescuing HDAC3’s enzymatic activity, the kinase activity of IPMK is required for HDAC3 activation.

**Figure 2:**
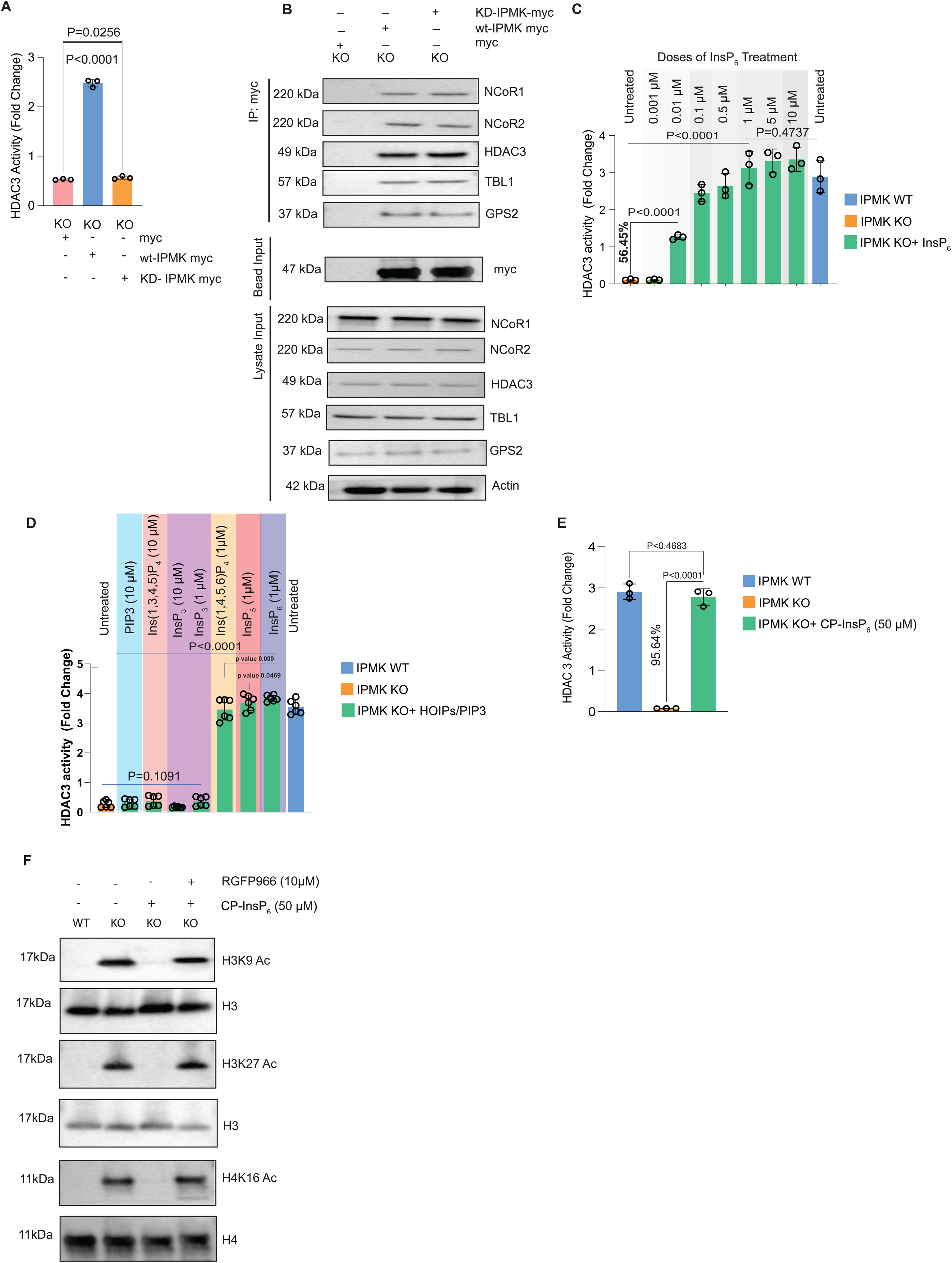
Kinase activity of IPMK is essential for activating HDAC3. **(A, B)** *IPMK*-Knockout (KO) HCT116 cells transiently overexpressed with *myc-control*, *WT-IPMK-*myc, and *KD-IPMK-*myc plasmid. Results demonstrated that *myc-control* and *IPMK-KD-*myc *were* unable to rescue HDAC3 activity while wt-IPMK-myc rescued. Data was represented as a fold change compared to IgG. Results were representative of three independent experiments (n=3).**(B)** Immunoprecipitation of IPMK-myc followed by western blot of NCoR1,NCoR2,HDAC3, GPS2, TBL1 indicates that the interaction of HDAC3,TBL1, GPS2 with NCoR1/2 in *IPMK* KO HCT 116 cells was comparable in *KD-IPMK-*myc as compared to *WT-IPMK-*myc. Data represents three experimental replicates (n=3). **(C)** To analyze the role of InsP_6_ in activating HDAC3 activity *in vitro*, endogenous HDAC3 were immunopurified from Wild type HCT116 (WT) and *IPMK-* Knock out HCT 116 (KO) cells followed by dose-dependent (0.001 µM, 0.01 µM, 0.1 µM, 0.5 µM, 1 µM, 5 µM and 10 µM) *in vitro* InsP_6_ treatment. Data was represented as a fold change compared to respective IgG. Results were representative of three independent experiments (n=3). 54.45% increase of HDAC3 activity after 0.01 µM of InsP_6_ treatment in IPMK KO cells. **(D)** To analyze the impact of other HOIPs and PIP3 in HDAC3 activity, endogenous HDAC3 were immunopurified from Wild Type HCT116 (WT) and *IPMK-* Knock out HCT 116 (KO) cells followed by *in vitro* 10µM treatment with PIP3, InsP_3_, Ins (1,3,4,5)P_4_ and 1µM of InsP_3_, Ins (1,4,5,6)P_4_, InsP5 and InsP_6_. Data was represented as a fold change compared to IgG, following blank subtraction and normalization. Results were representative of three independent experiments (n=3). **(E)** 24-hour treatment with 50 µM cell-permeable InsP_6_ (CP-InsP_6_) in IPMK-knockout (KO) HCT116 cells significantly restored HDAC3 activity comparable to untreated wild-type (WT) HCT116 cells. The results showed that 95.64% of HDAC3 activity was rescued in IPMK-KO cells after 24 hr of 50 µM CP-InsP_6_ treatment, relative to IPMK KO cells. Data was represented as a fold change compared to IgG. Results were representative of three independent experiments (n=3). **(F)** Cell-permeable InsP_6_ (CP-InsP_6_) treatment at a dose of 50 µM for 24 hr successfully rescued H3K9, H3K27 and H4K16 acetylation as comparable with untreated WT cells which HDAC3 inhibitor RGFP966 treatment abrogated. Data represents three experimental replicates (n=3).

Next, we tested the possibility of rescuing HDAC3’s enzymatic activity with InsP_4_, InsP_5_, and InsP_6_ *in vitro* test tubes after immunopurification of HDAC3 from *IPMK* KO cells. Biochemical analysis revealed that InsP_6_, in a dose-dependent manner, rescued the enzymatic activity of HDAC3 **(Fig 2 C)**. While 10 nM (0.01μM) InsP_6_ restored >50% of HDAC3 activity, 1 μM InsP_6_ [InsP (1,2,3,4,5,6) P_6_] fully restored activity comparable to the WT **(Fig 2 C)**. In WT cells, 5 or 10 μM InsP_6_ did not further increase HDAC3 activity, suggesting InsP_6_ saturation and a threshold effect at 1μM **(Fig 2B)**. Although InsP_4_ [InsP (1,4,5,6) P_4_] and InsP_5_ [InsP (1,3,4,5,6) P_5_] also restored HDAC3’s activity at 1 μM **(Fig 2 D)**, InsP_6_ showed the highest effect compared to all other HOIPs. Furthermore, while 10 nM of InsP_5_ and InsP_6_ both restored around 50% of HDAC3 activity compared to WT cells **(Extended Fig 2 A)**, InsP_4_ (10 nM) did not have any effect on HDAC3 activation and was similar to the KO cells. Interestingly, a second InsP_4_ isomer [InsP (1,3,4,5) P_4_] and InsP_3_ [InsP (1,4,5) P_3_] completely failed to rescue HDAC3 activity, even at 10 μM dose **(Fig 2 D)**. While Wu et al.’s^16^ previous study indicated InsP_3_ activating HDAC3, our experiments showed that InsP_3_ could not influence HDAC3 activity at either 1 μM or 10 μM concentrations **(Fig 2 D)**. This finding is supported by previous crystal structure analysis^6^, indicating the necessity of a phosphate group at the 6^th^ position of higher order inositolphosphate (HOIP) activating HDAC3. This crucial phosphate is missing in both InsP_3_ and the InsP_4_ isomer [InsP(1,3,4,5)P_4_], explaining their inability to activate HDAC3 in our study. As IPMK is a physiological PI3 kinase ^13^that also generates PIP3, we further tested the role of PIP3 in rescuing HDAC3 activity when purified from *IPMK* KO cells. We found that even at 10 μM, PIP3 failed to restore HDAC3 activity and was similar to the *IPMK* KO **(Fig 2 D)**. Based on the dose-dependent study **(Fig 2 C, D)** and previous research demonstrating InsP_6_’s health benefits^11^, we opted to utilize InsP_6_ for our subsequent experiments.

Next, we explored the possibilities of rescuing HDAC3 activity *in vivo* by treating cells with cell-permeable InsP_6_ (cell-permeable InsP_4_ and InsP_5_ have not been successfully synthesized). InsP_6_ exhibits poor cell permeability in cultured cells as cations present in cell culture media (such as calcium, magnesium, sodium, and potassium) may chelate InsP_6_, diminishing its availability for cellular uptake. Consequently, experiments often require the use of high, non-physiological concentrations of InsP_6_ to achieve desired effects. To overcome this, we used a cell-permeable InsP_6_ precursor molecule, where the negative charges of InsP_6_ are temporarily masked with biolabile groups, allowing it to pass through the cell membrane and then releasing active InsP_6_ after removal of the biolabile groups by enzymatic digestion^17^. We treated *IPMK* KO cells with this cell-permeable InsP_6_ and measured HDAC3 activity. Remarkably, 24 hours of cell-permeable InsP_6_ treatment rescued HDAC3 activity in *IPMK* KO cells to levels comparable to those of WT cells **(Fig 2 E)**, followed by a substantial decrease in histone acetylation, which was also comparable to the WT **(Fig 2 F)**. However, CP-InsP_6_ did not exert any effect when treated to wild type cells **(Extended Fig 2B)**. Notably, the treatment of HDAC3 inhibitor RGFP966 inhibited the rescue effect of cell-permeable InsP_6_ **(Fig 2 F)**, indicating that InsP_6_ deacetylates acetylated H3K9/27 and H4K16 by activating HDAC3.

Lastly, we evaluated the ability of CP-InsP6 to enhance intracellular InsP6 levels. Following 24 hours of treatment with 50 μM CP-InsP6, mass spectrometric analysis revealed a significant increase in intracellular InsP6 compared to wild-type cells (Extended Fig 2C). We also observed a parallel rise in InsP5 and InsP4 levels, suggesting dephosphorylation of InsP6 by phosphatases into lower-order inositol phosphates. Nevertheless, InsP6 remained the most abundant among the detected species.

### *IPMK* deletion disrupts HDAC3’s interaction with the DAD domain of its corepressor proteins, diminishing the activity of chromatin-bound HDAC3

Next, we wondered about the mechanism underlying the loss of HDAC3’s enzymatic activity in *IPMK*-deleted cells. As mentioned earlier, the HDAC3 protein level remains unaltered in *IPMK* KO cells **(Extended Fig 1B)**; hence, we wondered whether *IPMK* deletion disrupts HDAC3 binding to its corepressor protein, which is essential for HDAC3 function^1^. Immunoaffinity purification **(Extended Fig 3A)** and proximity ligation assay (PLA) **(Fig 3A, B)** demonstrated that *IPMK* deletion did not disrupt the association between HDAC3 and its corepressor protein, maintaining the overall integrity of the complex. While the SANT domain of the corepressor protein maintains the integrity of the HDAC3:corepressor complex^1,9^, the corepressor’s DAD domain binding to HDAC3 induces structural changes^1,9^,that are crucial for HDAC3’s enzymatic activity, although it is not necessary for preserving the complex’s integrity **(Fig 3C)**. Given this, it’s plausible that *IPMK* deletion specifically compromised the interaction between the DAD domain and HDAC3, thereby hindering HDAC3’s enzymatic activity without disrupting the overall complex integrity. To investigate whether *IPMK* deletion interferes with DAD domain interaction with HDAC3, we overexpressed DAD-Flag in *IPMK* knockout cells and subsequently performed a pull-down assay to isolate endogenous HDAC3. DAD-Flag pulled down endogenous HDAC3 from WT cells but, interestingly, failed when pulled down from *IPMK* KO cells **(Fig 3 D)**. Notably, the impaired DAD binding to HDAC3 observed in *IPMK* knockout cells was successfully rescued by treating the cells with cell-permeable InsP_6_ **(Fig 3 E)**. This observation aligns with previous crystal structure analyses, which revealed HOIP’s role as a molecular “glue” facilitating the interaction between the DAD domain of the corepressor protein and HDAC3^1,6,9^.

**Figure 3:**
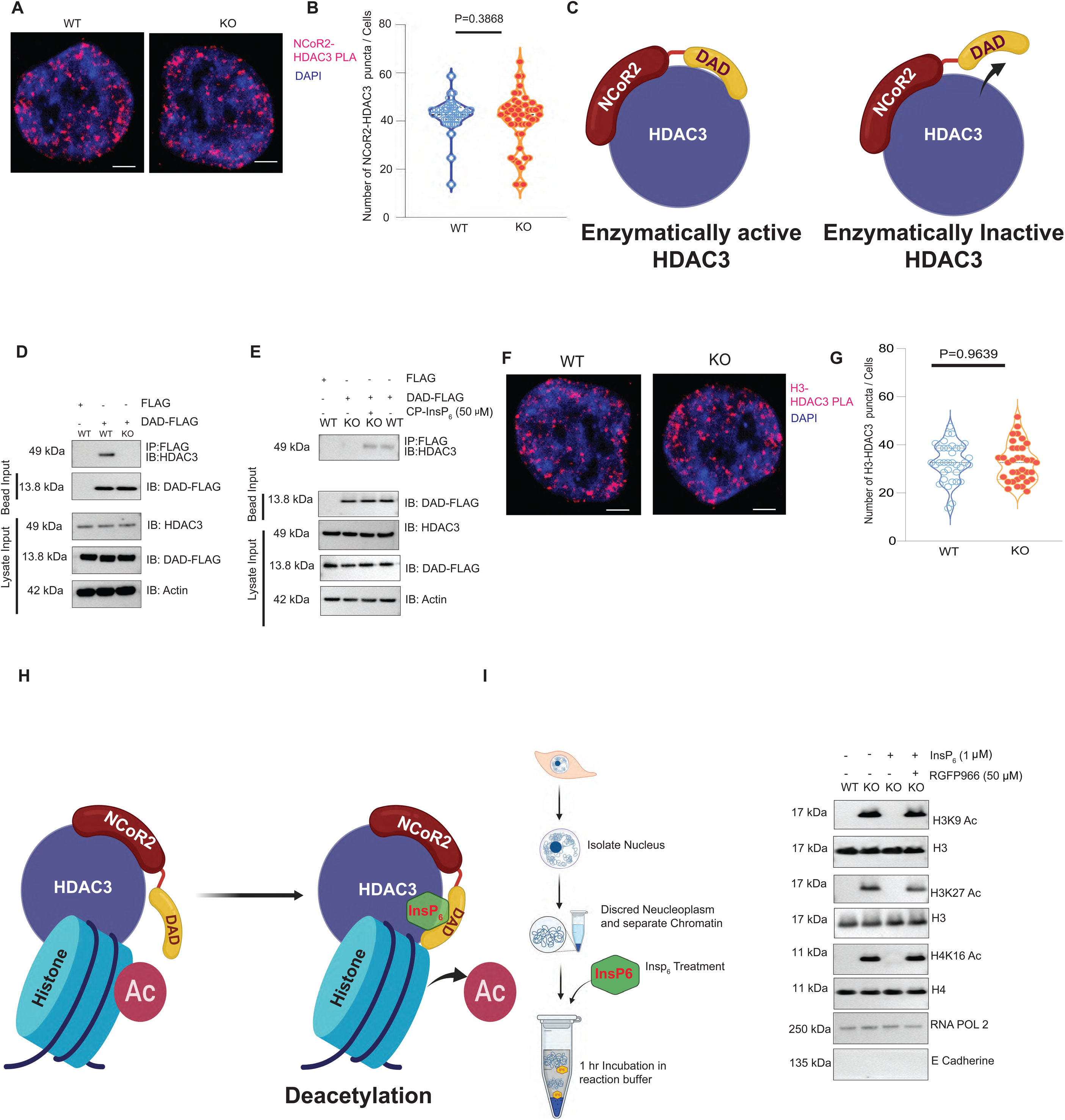
Mechanism of IPMK mediated HDAC3 activation *in vivo* and cell free chromatin fraction: **(A,B)** Proximity ligation assay using anti-HDAC3 and anti-NCoR2 antibody showed comparable amount of puncta (RED) in the WT and *IPMK* KO HCT 116 cells. Images were representative of three individual experiments (n=50). **(C)** Pictorial representation of NCoR2 protein domains, which includes the SANT1, SANT2 and DAD domains. The model represents that interaction of DAD of NCoR2 is crucial for the enzymatic activity of HDAC3, but does not influence HDAC3: corepressor complex integrity. **(D)** WT and *IPMK*-Knockout (KO) HCT116 cells transiently overexpressed with FLAG-control, and DAD-FLAG plasmid. FLAG was immunoprecipitated by WT and IPMK KO HCT 116 cells, followed by immunoblotting against anti-HDAC3 antibodies. Results demonstrated that DAD-FLAG failed to bind with endogenous HDAC3 when over expressed in *IPMK* KO cells while DAD-FLAG overexpressed in WT HCT 116 cells successfully to binds with endogenous HDAC3. Lysate control and bead inputs were represented. The result was representative of three individual experiments (n=3). **(E)** Treatment of *IPMK*-Knockout (KO) HCT116 cells with cell permeable InsP_6_ (CP-InsP_6_) at a dose of 50 µM for 24 hr,restored Flag-DAD binding to endogenous HDAC3. The result was representative of three individual experiments (n=3). **(F, G)** Proximity ligation assay with anti-HDAC3 and anti-H3 antibody showed comparable amount of puncta (RED) in the WT and *IPMK* KO HCT 116 cells. Results were representative of three individual experiments. Scale bar 5 μM (n=35). **(H)** Picture represents that HDAC3:NCoR2 complex binds to chromatin independent of InsP_6_; however, remains enzymatically inactive because DAD domain does not interact with HDAC3; presence of InsP_6_ drives DAD domain HDAC3 interaction activating HDAC3’s deacetylase activity. **(I)** Schematic diagram of chromatin purification and *in vitro* InsP_6_ treatment. InsP_6_ treatment to cell free chromatin fraction isolated from IPMK KO cells repressed H3k9/27 and H4k16 acetylation which HDAC3 inhibitor RGFP966 treatment abrogated. Immunoblotting against anti-RNA POL 2 (chromatin) and E Cadherin (cytoplasmic and cell membrane) antibody used as cell fractionation control. Data represents three experimental replicates (n=3).

Given that HDAC3’s chromatin recruitment is crucial for HDAC3-mediated histone deacetylation next, we wondered whether *IPMK* deletion impaired HDAC3’s chromatin recruitment. Using HDAC3-to-histone H3 PLAs, or immunopurification of H3 or H4 chromatin fraction followed by HDAC3 western blot, we found a comparable amount of HDAC3 chromatin recruitment in *IPMK* KO and WT cells **( Fig 3 F, G, Extended Fig 3B)**, indicating that *IPMK* deletion did not impair HDAC3’s chromatin recruitment. Our data collectively support a functional model where HDAC3 recruitment to chromatin and its binding to the corepressor complex occur independently of IPMK. However, InsP_6_ is crucial for the DAD domain’s interaction with HDAC3, which triggers its enzymatic activity and subsequent histone deacetylation **(Fig 3H)**.

To investigate whether InsP_6_ serves as the final rate-limiting step in HDAC3-mediated histone deacetylation, we employed a simplified cell-free system. We treated cell-free, chromatin fraction in reaction buffer from *IPMK* knockout cells with InsP_6_ for one hour, a context where any observed effects would likely result from InsP_6_’s direct action on chromatin-bound HDAC3, independent of complex multistep processes of HDAC3 chromatin recruitment involving other proteins. Strikingly, InsP_6_ treatment fully suppressed histone acetylation in *IPMK* KO nuclear lysate **(Fig 3I)**, restoring it to wild-type levels. The specificity of this effect was confirmed by co-treatment with the HDAC3-selective inhibitor RGFP966, which negated InsP_6_’s impact **(Fig 3I)**. However, the overall integrity of the HDAC3 chromatin protein complex (including NCoR1/2, TBL1, GPS2)^18^, remains intact in *IPMK* KO cells and is comparable to the WT **(Extended Fig 3C)**. These results provide strong evidence that InsP_6_ directly activates chromatin-bound HDAC3 by recruiting the DAD domain of the corepressor to HDAC3, functioning as the critical final step in the histone deacetylation process.

IPMK generates InsP5, while IPPK produces InsP_6_. Given that IPMK interacts with HDAC3, we examined whether IPPK also associates with HDAC3, potentially contributing to a localized HOIP signaling event within the HDAC3 complex. Our biochemical and microscopy analyses revealed that IPPK indeed interacts with nuclear HDAC3 (Extended Fig. 3II, A, B).

### Disruption of the IPMK-HDAC3 epigenetic axis hypertranscribes MMP gene family

We aimed to investigate the role of the IPMK-HDAC3 epigenetic axis in transcriptional regulation. Our bulk RNA sequencing analysis of *IPMK*-deleted HCT116 cells revealed significant transcriptional changes compared to wild-type cells, with 4,014 genes upregulated and 3,271 downregulated **(Fig 4A)**. Pathway analysis indicated that the genes related to the extracellular matrix degradation, which includes the collagenase gene family, were notably upregulated in *IPMK* KO cells **(Fig 4B)**. Specifically, archetypal matrix metalloproteinases (MMPs), including MMP1, MMP10, MMP12, and MMP13, were significantly upregulated in the RNA-seq data **(Fig 4A)**. We validated these findings at the protein level using western blot analysis for MMP1, MMP10, and MMP13, confirming the RNA-seq results **(Fig 4C)**. Due to a lack of high-quality antibodies, we were unable to test MMP12.

**Figure 4:**
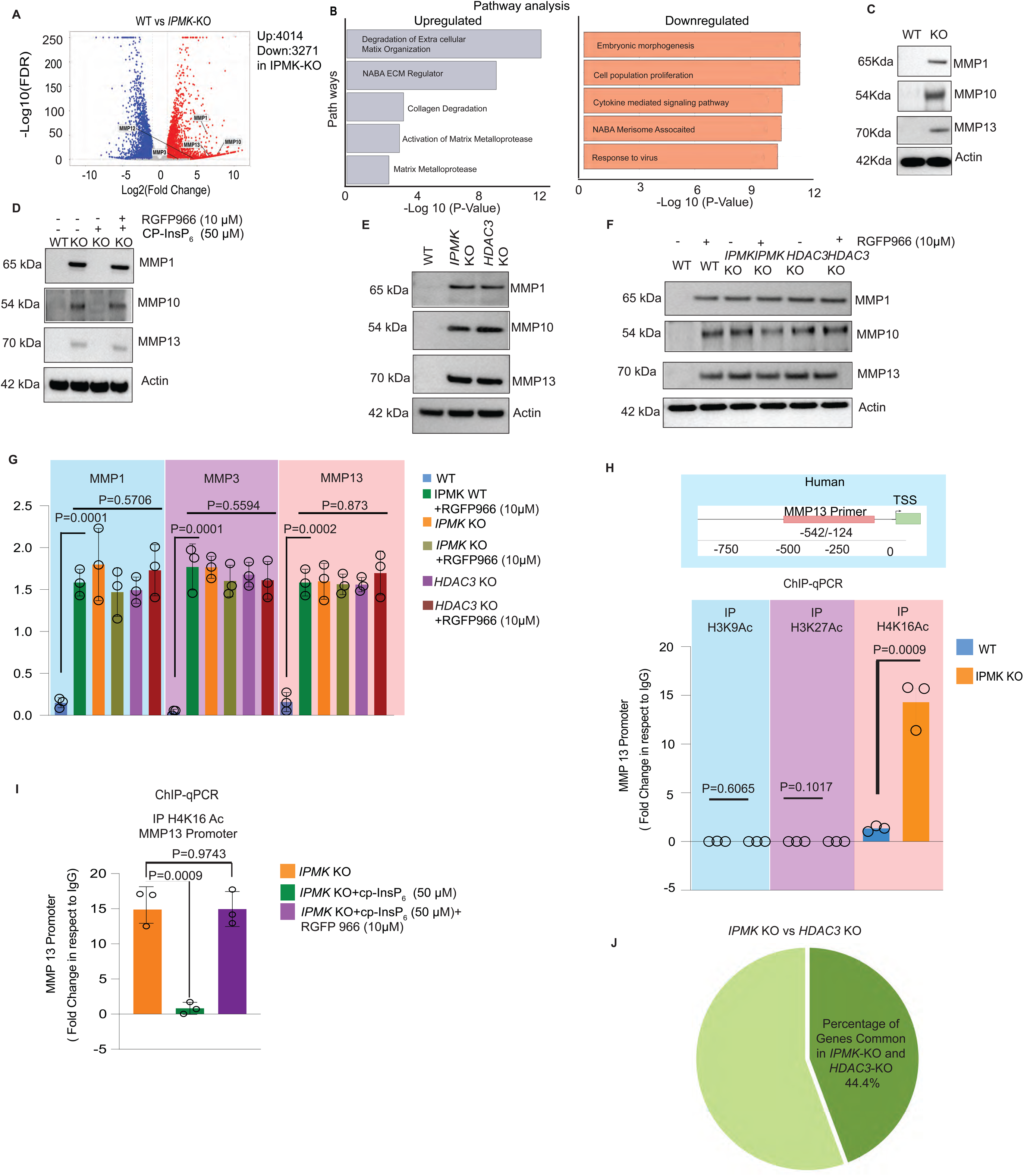
Deletion of *IPMK* stimulates hypertranscription of MMP gene family. **(A)** Volcano plot (Log 10[*FDR*] vs. Log 2[Fold change]) displays differentially expressed genes after *IPMK* deletion in HCT116 cells (*IPMK-KO*) as compared to WT-HCT116 cells (*WT*). Red dots represent gene expression that was upregulated, while blue dots represent gene expression that was downregulated in *IPMK* KO cells. Y-axis denotes – Log10 FDR Value, while the X-axis denotes the Log2 fold change value. The result was representative of three individual experiments (n=3). **(B)** Pathway analysis of RNA-seq data identified common genes that are upregulated and down regulated following *IPMK* deletion in HCT 116 (KO) cells. **(C)** Western blot analysis showed MMP1, 10, and 13 markedly increased in *IPMK* KO HCT116 cells. (n=3) **(D)** Western blot of MMP1, 10 and 13 from IPMK WT and KO cells. 24 hr of 50 µM CP-InsP_6_ treatment to KO cells successfully rescued MMP1, MMP10 and MMP 13 expression while co treatment with 10 µM RGFP966 abrogated InsP_6_’s rescue effect. Actin used as loading control. Data represents three experimental replicates (n=3). **(E)** Lysate isolated from WT, *IPMK*-KO, and *HDAC3* KO HCT 116 cells were western blotted against anti-MMP1, anti-MMP10 and anti-MMP13 antibody. Result demonstrated as compared to WT HCT 116 cells (WT) MMP1, MMP10 and MMP 13 expression was increased in *HDAC3* KO HCT 116 cells (*HDAC3* KO) which is comparable with *IPMK* KO HCT116 (*IPMK* KO) cells. Actin used as loading control. Data represents three experimental replicates (n=3). **(F)** Western blot analysis revealed that the deletion of *IPMK or HDAC3* led to profound increased MMP1, MMP10 and MMP13 expression as compared to WT HCT116 cells. An 8-hour treatment with the HDAC3-specific inhibitor RGFP966 at a dose of 10 µM in HCT116 WT, *IPMK* KO, and *HDAC3* KO cells showed no further increase in MMP1, MMP10, MMP13 expression compared to the untreated *IPMK* KO and *HDAC3* KO HCT116 cells. Actin used as loading control. Data represents three experimental replicates (n=3). **(G)** Representative densitometric analysis. Data represents three experimental replicates (n=3). **(H)** Pictorial representation showing MMP13 primer locations on the human genome. Chromatin immunoprecipitation against acetylated H3K9, H3K27 and H4K16 followed by qPCR analysis using primers for MMP13 promoter. Results depicted that *IPMK* deletion selectively induced promoter enrichment of acetylated H4k16, but not for acetylated H3k9/27. **(I)** MMP13 promoter enrichment was significantly rescued in *IPMK*-KO cells after 24 hr 50 µM cell permeable InsP_6_ (CP-InsP_6_) treatment. Co-treatment of 10 µM RGFP966 with 50 µM cell permeable InsP_6_ (CP-InsP_6_) abrogated InsP_6_’s rescue effect (n=3). **(J)** Overlap of differentially expressed RNA seq data compared with *IPMK* KO and *HDAC3* KO cells.

Given that IPMK is recognized as a risk gene for Inflammatory bowel disease (IBD)^13^, we also analyzed transcriptomic data from human patients using the Inflammatory Bowel Disease Transcriptome and Metatranscriptome Meta-Analysis (IBDTaMMA^19^, https://ibd-meta-analysis.herokuapp.com/)^19^ **(Extended Fig 4 A)**. This analysis revealed significant transcriptional upregulation of MMP1, MMP3, MMP10, MMP12, and MMP13, in IBD patient cells similar to the findings in *IPMK* KO HCT116-intestinal cells. Furthermore, separate analyses of Crohn’s disease (CD) and ulcerative colitis (UC) patients revealed a striking increase in major MMPs—such as MMP3, MMP13, and MMP10—that were consistently elevated in patient tissues and similarly upregulated in IPMK-depleted cells (Extended Fig 4 II A,B).

Interestingly, *IPMK* KO cells rescued with *wild-type IPMK* showed a reduction in MMP expression, whereas cells with kinase-dead IPMK did not exhibit this effect **(Extended Fig. 4B)**. Importantly, cell-permeable InsP_6_ treatment to *IPMK* KO cells suppressed MMP expression as well, making it comparable to that of the WT cells **(Fig 4D)**, indicating that the loss of IPMK’s inositol kinase activity upregulates MMP gene expression. To determine whether InsP_6_ treatment repressed MMP gene expression by activating HDAC3, we co-treated HDAC3 inhibitor RGFP966, which abrogated the rescue effect of InsP_6_ **(Fig 4D)**.

Moreover, in the *HDAC3* KO cells, MMP genes **(Extended Fig 4C)** and protein expression were markedly increased **(Fig 4E)** and comparable to the *IPMK* KO cells **(Fig 4E)**. In our second approach to determine whether identified MMP genes are regulated via the IPMK-HDAC3 epigenetic axis, we examined the effects of the HDAC3 inhibitor RGFP966 on WT, *IPMK* KO, and *HDAC3* KO cells. We hypothesized that RGFP966 would increase MMP expression in WT cells but not in *IPMK or HDAC3* KO cells, as these cells lack HDAC3 activity. However, if MMP expression in *IPMK* KO cells was influenced by pathways independent of HDAC3, RGFP966 treatment might further increase MMP expression. As anticipated, RGFP966 significantly increased MMP expression in WT cells but failed to induce further expression in *IPMK and HDAC3* KO cells **(Fig 4F, G)**. Taken together, the results confirm that the disruption of the IPMK-HDAC3 epigenetic axis is responsible for the transcriptional upregulation of MMPs, ruling out the involvement of HDAC3-independent pathways in this process.

Next, we examined whether the promoters of matrix metalloproteinases (MMPs) exhibit enrichment of histone acetylation. We selected MMP13 as a representative example for this analysis. Our biochemical studies confirmed that *IPMK* deletion selectively upregulates H3K9/27 and H4K16 acetylation at the protein level, an effect also observed in *HDAC3* KO cells and those treated with HDAC3 inhibitors **(Fig 1F)**. Intriguingly, ChIP-qPCR targeting the MMP13 promoter region **(Fig 4H)** revealed a selective enrichment of acetylated H4K16 in the promoters of MMP13 in *IPMK* KO cells, while H3K9/27 acetylation showed no significant change **(Fig 4 H)**. The ChIP data also indicated an enrichment of HDAC3 as tested using MMP13 promoter regions **(Extended Fig 4 D)**.

A particularly fascinating finding emerged when we treated *IPMK KO* cells with cell-permeable InsP_6_. This treatment substantially reduced the enrichment of acetylated H4K16 at MMP promoters **(Fig 4I)**. However, this effect was nullified when cells were co-treated with the HDAC3 inhibitor RGFP966 **(Fig 4I)**. Notably, InsP_6_ treatment did not affect HDAC3 chromatin recruitment **(Extended Fig 4 D)**.

Given the known upregulation of MMP3 in IBD^20^, we revisited our *IPMK* KO HCT116 RNA-seq data to understand the MMP3 log fold change. Although we found a 2.5-Log 2-fold increase in MMP3 expression in HCT116 cells, the p-value of 0.3 exceeded our significance threshold of p<0.05. Raw data analysis revealed zero reads in all three WT samples and a maximum of two reads in KO samples for MMP3, likely contributing to the lack of statistical significance. While deeper sequencing might yield statistically significant results, our qPCR **(Extended Fig 4E)** and protein-level data **(Extended Fig 4 F - H)** clearly demonstrated a substantial increase in MMP3 levels in both *IPMK* and *HDAC3* KO cells, which was reversed by InsP_6_ treatment **(Extended Fig 4I)**. Furthermore, studies using HDAC3 inhibitors and knockout models confirmed that disrupting the IPMK-HDAC3 epigenetic axis substantially elevated MMP3 transcriptional expression, providing additional validation of our findings.

Finally, to assess the overlap between the global transcripts regulated by HDAC3 and IPMK, we first identified differentially expressed genes in *HDAC3* KO cells and compared this dataset with the expression profile of *IPMK* KO cells. The results showed that approximately 44% of HDAC3-regulated genes were also affected in *IPMK* KO cells **(Fig 4J)**. Importantly, HDAC3 exhibits a dual regulatory role in transcription: it functions not only through its deacetylase activity but also as a cotranscriptional activator independent of this enzymatic function^16^. This dual functionality likely explains why certain genes are affected by *HDAC3* deletion but remain unaltered in *IPMK* KO cells. Specifically, the genes impacted in *HDAC3* KO cells but not in *IPMK* KO cells may be influenced by HDAC3’s non-enzymatic roles, such as its function as a cotranscriptional activator.

### *IPMK* deletion is associated with increased cell and intestinal permeability in mice

The upregulation of MMP is associated with compromised intestinal barrier function by disrupting tight junctions, leading to a “leaky gut”, which is commonly observed in IBD patients^12^. Given these associations, we investigated the consequences of *IPMK* deletion stimulating increased MMP production in cell permeability. To assess that, we employed a transwell assay using Caco2 cells, which is classically used for *in vitro* cell permeability assay^21^. We cultured these cells in a confluent layer on a trans well insert and treated them with 4kDa FITC-Dextran. Cell permeability would increase the leakage of 4kDa FITC-Dextran from the upper cell chamber into the bottom chamber **(Fig 5 A)**. Notably, conditioned media from *IPMK* KO or *HDAC3* KO cells elevated Caco2 cell permeability **(Fig 5 B)**, while media transferred from WT cells failed to induce cell permeability. The observed effect may be attributed to the secretory nature of MMPs. In *IPMK* KO HCT116 cells, the conditioned media contained elevated levels of secreted MMPs (Extended Fig 5 II A). When this MMP-enriched media was transferred to wild-type Caco2 cells, it increased cellular permeability by disrupting the cell-cell junction. This hypothesis was corroborated by the western blot study that identified the loss of tight junction proteins ZO-1 and Occludin in CaCo2 cells treated with either *IPMK* KO cell-conditioned media **(Extended Fig 5A)**. Remarkably, *IPMK or HDAC3* KO cells pretreated with Pan-MMP inhibitor (Ilomastat) followed by media transfer completely blocked the Caco2 cell permeability **(Fig 5 B)**, confirming that the effect was due to MMP in the conditioned media.

**Figure 5:**
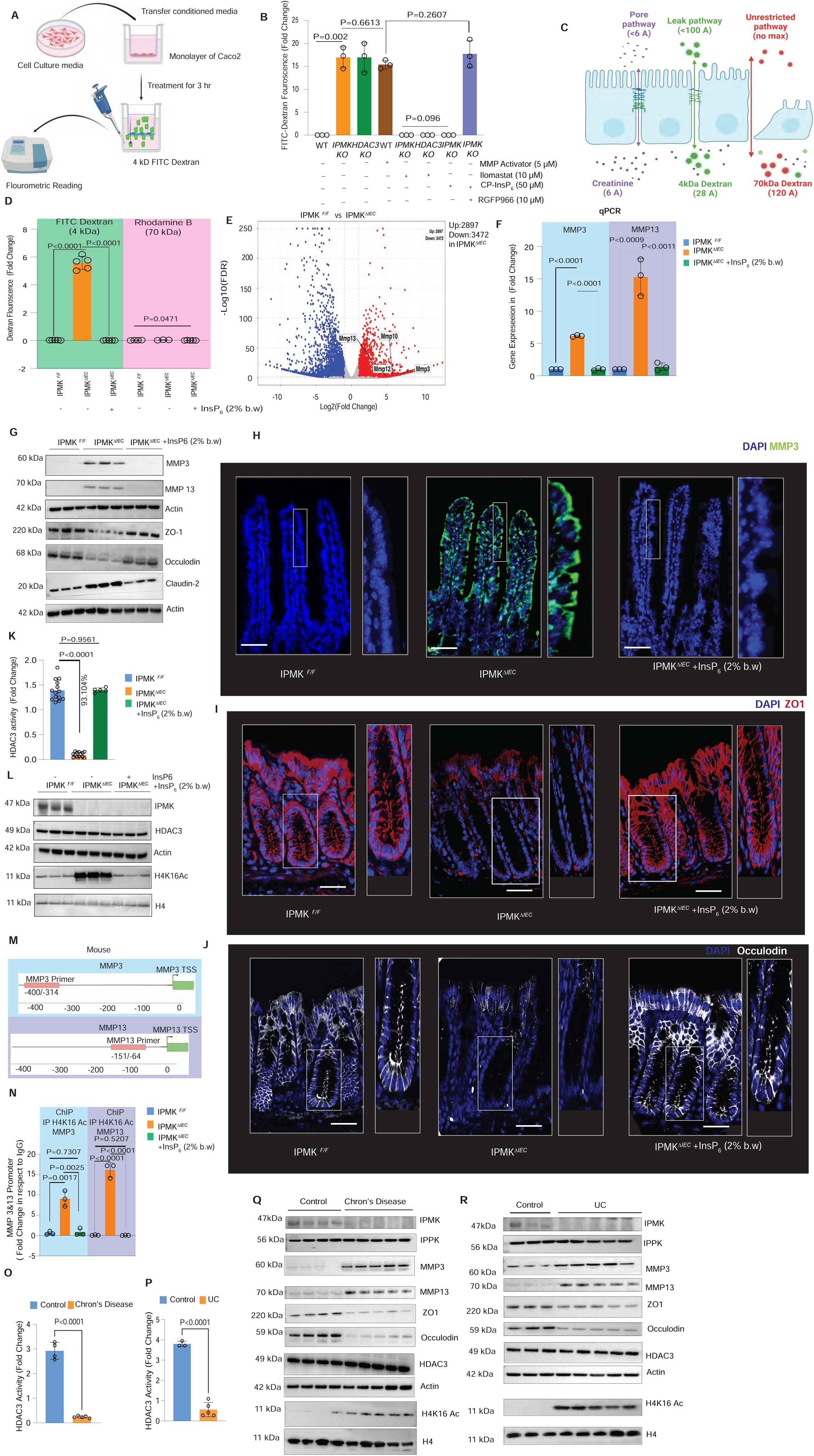
*IPMK* deletion markedly increased cell and intestinal permeability in mice. **(A)**Schematic diagram of transwell cell permeability assay. **(B)** Conditioned media transferred from *IPMK and HDAC3* KO cells to the Caco2 (WT) monolayer elevated cell permeability. Treatment of MMP activator (p-Aminophenylmercuric acetate) to WT HCT16 cell followed by conditioned media transfer also increased Caco2 cell permeability. Treatment of HDAC3 inhibitor RGFP966 to WT HCT16 cell followed by conditioned media transfer also increased Caco2 cell permeability. Treatment of CP-InsP_6_ to *IPMK* KO and *HDAC3* KO cells followed by conditioned media transfer to CaCo2 failed to induce cell permeability. RGFP966 co-treatment with InsP_6_ to *IPMK* KO cells followed by conditioned media transfer increased CaCo2 cell permeability Result was representative of three independent experiments. (n=3). **(C)** Pictorial representation of Pore, Leak and Unrestricted intestinal permeability pathway. Pore pathway does not allow 4kDa Dextran while Leak pathway allows 4kDa dextran permeability but not 70 kDa. Unrestricted pathway is activated due to cell death or cell damage allows both 4kDa and 70 kDa Dextran. **(D)** Intestinal permeability assay indicated a significant increase in 4 kDa FITC Dextran fluorescence (not 70 kDa Rhodamine) in IPMK^Δ^*^IEC^* mice serum compared to the IPMK *^F/F^* Interestingly, 3 days pretreatment of 2% body weight of oral InsP_6_ to IPMK^Δ^*^IEC^* reduced intestinal 4 kDa FITC Dextran leak. After 4 hr of fasting a single dose of 4 kDa FITC-Dextran was gavaged at 0.6mg/gm body weight. The untreated control group received an equal amount of PBS. Fluorescence intensity of plasma samples was measured (excitation 485 nm/emission 535 nm) after 4 hours of gavage. Data were expressed as fold change relative to control mice serum, which received only PBS gavaged mice without FITC or Rhodamine Dextran. Result was representative of three independent experiments (n=5). **(E)** Volcano plot (Log 10[*FDR*] vs. Log 2[Fold change]) displays differentially expressed genes in IPMK^Δ^*^IEC^* as compared with IPMK*^F/F^* mice. Red dots represent gene expression that was upregulated in IPMK^Δ^*^IEC^*, while blue dots represent gene expression that was downregulated. The Y-axis denotes –the Log10 FDR Value, while the X-axis denotes the Log2 fold change value. The result was representative of three individual experiments (n=3). **(F)** qPCR analysis of mRNA isolated from intestinal epithelial cells of IPMK*^F/F^* and IPMK^Δ^*^IEC^* cells indicated increased MMP3 and MMP13 gene transcription. 18s used as loading control. Results represented three experimental replicates (n=3). **(G)** Western blot from intestinal crypts (Ileum) showed increased MMP3 and 13 expression in IPMK^Δ^*^IEC^* mice while 72 hr 2% b.w InsP_6_ treatment reduced it. ZO-1, occludin level decreased with Caludin 2 level increased in IPMK^Δ^*^IEC^* mice; however, the level was restored by oral InsP6. The result was representative of three independent experiments (n=3). **(H)** Immunofluorescence microscopy showed increased MMP3 expression (Green) in ileum IPMK^Δ^*^IEC^* as compared to IPMK*^F/F^* .DNA (blue). 2% body weight (B.W) of InsP_6_ treatment for 72hr to IPMK^Δ^*^IEC^* completely reduced MMP3 expression. Scale bar represents 50 μm.Data was representative of three individual experiments. (n=3) (Total number of images=30). **(I,J)** Showing confocal images of colon ZO-1 (red pseudo color) and Occludin (white pseudo color) showing loss of its expression in IPMK^Δ^*^IEC^* and restored by InsP_6_. Scale bar 50 μm.**(K)** HDAC3 activity was diminished in intestinal epithelial cells isolated from IPMK^ΔIEC^ compared with IPMK*^F/F^*. 2% body weight (B.W) of InsP_6_ treatment for 72hr to IPMK^Δ^*^IEC^* completely reduced HDAC3 activity. Data was represented as a fold change compared to IgG. Results are representative of a minimum of three individual experiments. (n=14 for IPMK*^F/F^*, n=14 for IPMK^Δ^*^IEC^*, n=5 for 2% body weight (B.W) of InsP_6_ treatment for 72hr to IPMK^Δ^*^IEC^*). **(L)** Western blot from intestinal crypts (Ileum) indicated elevated H4K16 acetylation in IPMK^ΔIEC^ mice which was reduced by pretreatment of 2% oral InsP_6_. Western blot analysis further showed HDAC3 protein levels were comparable in all untreated and untreated genotypes. Actin was used as a loading control. Results were representative of three individual experiments (n=3). **(M,)** Pictorial representation showing MMP3 and 13 primer locations on the mouse genome. **(N)** ChIP-qPCR analysis indicated MMP3/13 promoter enrichment of acetylated H4K16 when purified from IPMK^ΔIEC^ crypts (Ileum), which was diminished by pretreatment oral InsP_6_ (2%). Data represents three experimental replicates (n=3). **(O)** Reduced HDAC3 activity in Crohns patient tissue (Control, n=4, CD,n=5) and **(P)** Ulcerative colitis patient tissue (Control, n=3, UC,n=5)**. (Q)** and **(R)** showing loss of IPMK in CD and UC patient samples. IPPK level unchanged. MMP levels increased in diseased tissue. ZO-1 and Occludin reduced in CD and UC. H4K16acetylation induced in CD and UC.

Furthermore, *IPMK* KO cells treated with just cell permeable InsP_6_ followed by media transfer to Caco2 cells prevented the Caco2 cell permeability, but the RGFP966 co-treatment abrogated InsP_6_’s rescue effect **(Fig 5 B)**. This confirms that disruption of the IPMK-HDAC3 epigenetic axis leads to MMP upregulation and consequent increased cell permeability, effects successfully rescued by InsP_6_ treatment.

*In vivo,* mice studies further supported the cell-level findings. We created intestine-specific *IPMK* KO mice (IPMK^ΔIEC^) **(Extended Fig 5 B-E)** and tested gut permeability by orally gavaged mice with FITC-dextran (4kD) and Rhodamine-dextran (70 kD). The disruption of the tight junction only allows 4kD dextran but restricts 70 kD^22–24^, activating the leaky pathway, while significant destruction of the epithelial cell layer allows both 4kD and 70 kD dextran **(Fig 5 C)**. Notably, the IPMK^ΔIEC^ mouse showed a marked increase in the leaky pathway, allowing 4kD dextran but without allowing 70 kD to pass through **(Fig 5 D)**, indicating intestinal loss of *IPMK* selectively activates the leaky pathway expected due to loss of tight junction disruption.

To determine the hyper-transcription of MMPs in IPMK^ΔIEC^ mice, we flow purified **(Extended Fig 5 B-E)** pure intestinal epithelial cells (IEC) from IPMK^F/F^ and IPMK^ΔIEC^ mice followed by RNA-seq. Data indicated a profound increase in MMP transcription **(Fig 5 E),** also validated by qPCR, tested in two representative MMP genes (MMP3 and MMP13) **(Fig 5 F).** Pathway analysis using Metascape (Extended Fig. 5II G-I) and GSEA **(Extended Fig. 5II J)** confirmed enrichment of gene sets associated with MMP activity and extracellular matrix organization. Protein level analysis by performing western blots of crypts **(Fig 5G, Extended Fig 5 II B,C)** and immunohistochemistry **(Fig 5 H)** reconfirmed transcriptomic data showing elevated MMPs in IPMK^ΔIEC^ mice. Fascinatingly, oral administration of InsP_6_ solution normalized to pH 7.0, treated to IPMK^ΔIEC^ mice completely reduced MMP levels, making them comparable to those observed in IPMK*^F/F^* mice **(Fig 5 G,H, Extended Fig 5 II B,C).** Next, we studied the levels of major tight junction proteins like ZO-1, Occludin, and claudin 2. Notably, ZO-1 and Occludin levels were significantly reduced in IPMK^ΔIEC^ mice as tested in Western blot and IHC studies **(Fig 5 G,I,J, Extended Fig 5 II D,E)**. Interestingly, claudin 2 levels were rather increased in IPMK^ΔIEC^ **(Extended Fig 5 II F)**, which agrees with previous studies indicating increased claudin 2 is linked with elevated intestinal permeability^23^. Importantly, oral InsP6 treatment rescued all tight junction protein levels. Moreover, the HDAC3 activity assay indicated a marked decline in HDAC3 activation in IPMK^ΔIEC^ **(Fig 5 K)**. Notably, oral InsP_6_ treatment restored HDAC3 activity to the level of IPMK*^F/F^* mice **(Fig 5 K**. Protein level analysis showed that in IPMK^ΔIEC^ mice intestine H4K16 acetylation was markedly increased which oral InsP_6_ restored back to the IPMK*^F/F^* level; however, the IPMK protein was unaffected by InsP_6_ treatment **(Fig 5 L, Extended Fig 5 F)**, indicating oral InsP_6_ treatment alone could restore the epigenetic change of H4K16 acetylation in IPMK^ΔIEC^ mice in absence of IPMK **(Fig 5L).** The protein level epigenetic change was reconfirmed at the chromatin level by ChIP-qPCR analysis of MMP3, and 13, indicating in IPMK^ΔIEC^ mice marked increase of H4K16ac in MMP3, 13 promoters, which were reduced and reversed back to the level of IPMK*^F/F^* mice by oral InsP_6_ treatment **(Fig 5 M,N)**. As expected, MMP promoter recruitment of HDAC3 was evident in IPMK^ΔIEC^ mice and comparable to the IPMK*^F/F^* mice, and it was unaffected by oral InsP_6_ treatment **(Extended Fig 5G).** While InsP_6_ cellular uptake is poor in cell culture systems, in our animal studies, a 2% body weight, InsP_6_ treatment was sufficient to demonstrate its effects. This observation aligns with previous research showing intestinal absorption^25–27^ and therapeutic efficacy of oral InsP ^28^ in mice and rats.

To determine whether the above finding has any clinical significance in relation to IBD, we found that both CD and UC patient samples showed diminished HDAC3 activity **(Fig 5 O, P)**. Remarkably, IPMK protein levels were markedly reduced in both CD and UC patient samples **(Fig 5Q,R)**, but this did not affect IPPK protein levels, indicating specificity. Moreover, we found a loss of ZO-1 and Occludin levels in CD and UC patient samples, along with increased H4K16 acetylation **(Fig 5Q,R)**.

The collective findings suggest that the deletion of intestinal *IPMK* leads to a reduction in HDAC3 activity, which in turn promotes the expression of (MMPs). This phenomenon is evident in IBD patient samples, showing a pronounced loss of IPMK and its downstream effects, as observed in IPMK^ΔIEC^ mice. This cascade of events disrupts cell junctions, ultimately resulting in a leaky gut condition. Notably, oral administration of InsP_6_ effectively reversed leaky gut effects.

### In a mouse model of IBD, *IPMK* expression and HDAC3 activity are reduced, and oral InsP_6_ treatment restores intestinal permeability

We used a DSS (dextran sodium sulfate)-induced IBD mice model ^29^ to test the leaky gut, where, marked elevation in intestinal permeability for 4kD dextran was evident as early as 24h, not allowing 70kD **(Fig 6A)**. Remarkably, pre-treatment with oral InsP_6_ dosed at 2% of the mouse body weight 72h prior to the DSS treatment, prevented leaky gut comparable to the untreated mice **(Fig 6 A)**. Since InsP_6_’s bioavailability is modest^25^, we pretreated mice for 72 hours. Notably, the 24h of DSS treatment selectively reduced IPMK protein expression in the HOIP pathway **(Fig. 1 A)** while keeping IPPK protein expression unaltered (**Fig 6B, Extende Fig. 6A),** with a substantial loss of HDAC3 activity, which was restored in DSS-treated mice by oral InsP_6_ pre-treatment **(Fig 6C)**. Moreover, the western blot of colonic crypts indicated a substantial increase in MMP3 and 13 expressions after DSS treatment, which oral InsP_6_ pretreatment attenuated and restored to the level of untreated mice **(Fig 6D)**. We reconfirmed the western blot data using immunohistochemistry of MMP3 in the colon and even in the ileum, showing that both small and large intestines exhibited increased MMP3 expression **(Fig 6E-H)**.

**Figure 6:**
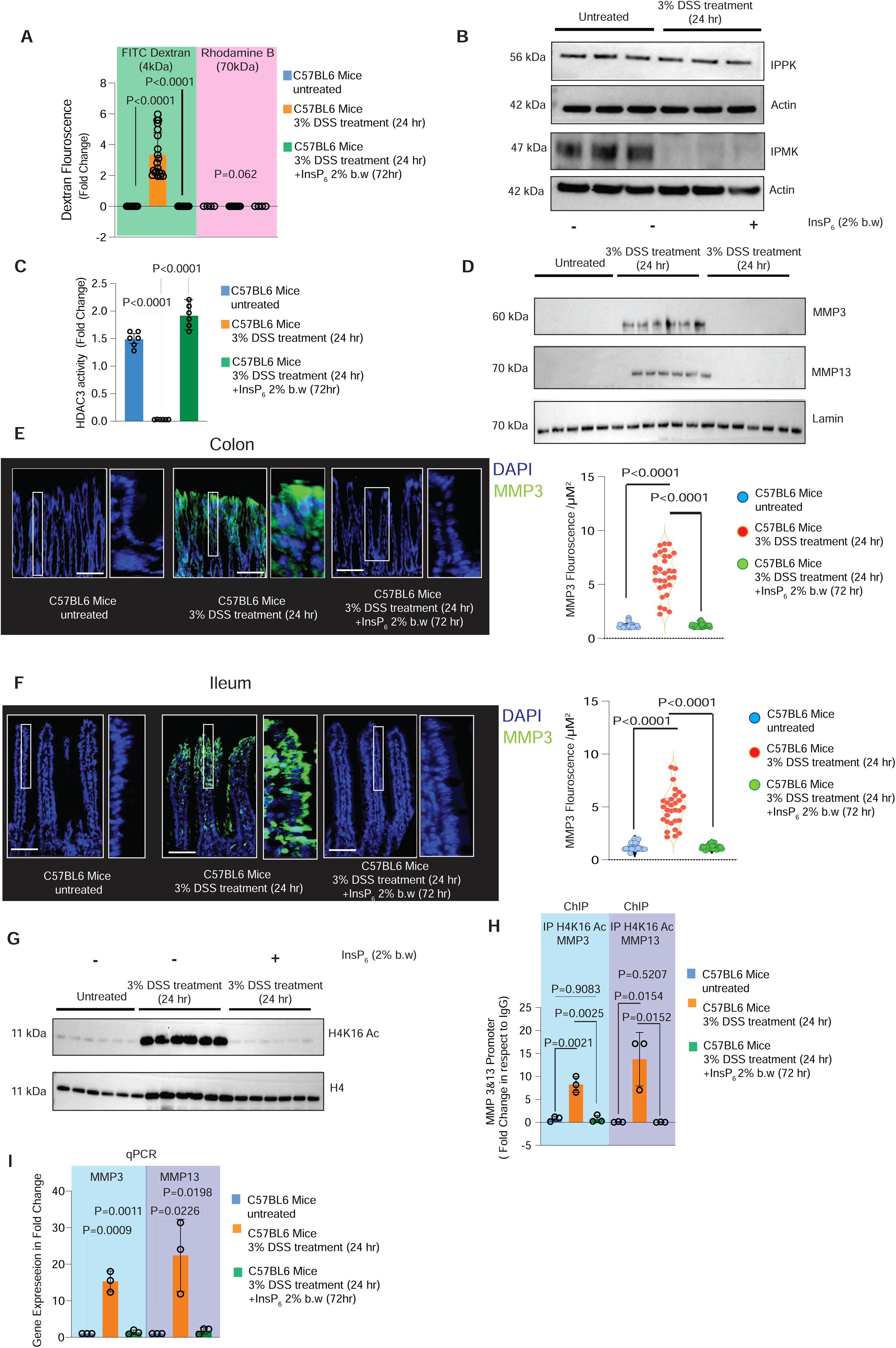
In the DSS-induced IBD mouse model, diminished IPMK expression and HDAC3 activity led to increased intestinal permeability, which was rescued by oral InsP_6_ treatment. **(A)** Intestinal permeability assay indicated a significant increase in 4 kDa FITC Dextran fluorescence in 24 hr 3% DSS treated mice serum without altering 70 kDa Dextran (Rhodamine B) as compared with untreated mice. Data were expressed as fold change relative to control mice serum, which received only PBS gavage without FITC or Rhodamine Dextran. (n=15). **(B)** Western blot from colonic crypt showed that 3% DSS treatment reduced IPMK expression while IPPK expression was unaltered. Actin was used as loading control. (n=3) **(C)** 3% DSS treatment (24 hrs) markedly reduced HDAC3 activity in colonic crypts which oral InsP_6_ treatment rescued. Result was representative of three independent experiments (n=6). **(D)** Western blot from colonic crypts indicated increased MMP3 and 13 expression after 3% of DSS treatment (24hrs) which pretreatment of oral InsP_6_ reduced. Actin was used as loading control. Data was representative of three individual experiments. (n=6) **(E)** Immunofluorescence microscopy shows increased MMP3 (Green) in 24 hr 3% DSS treated colon and **(F)** ileum as compared to untreated control. That was reduced by InsP_6_ pretreatment (72hr). Scale bar represents 50 μm. Data represents three experimental replicates (n=3). (Total number of images=30) **(G)** Western blot from colonic crypt showed that 3% DSS treatment induced H4k16 acetylation which InsP_6_ pretreatment (72hrs) reduced. Actin was used as loading control. Result was representative of three independent experiments (n=6). **(H)** ChIP-qPCR analysis indicated MMP3/13 promoter enrichment of acetylated H4K16 when purified from DSS (3%) treated colonic crypts, which was diminished by pretreatment oral InsP_6_ (2%). Data represents three experimental replicates (n=3). **(I)** qPCR analysis of MMP3 and 13 isolated from colonic crypt. 24 hrs of 3%DSS treated induced gene expression while 2% of InsP_6_ treatment reduced it. 18s used as loading control. Results represented three experimental replicates (n=3).

Though acute DSS treatment mostly shows ulcerative colitis-like phenotype in mice colon, the previous study also indicated its pathological effects in the ileum^30^; hence, we also included the ileum to show the molecular effect is unrestricted throughout the intestine after DSS treatment.

At the epigenetic level, DSS treatment markedly increased H4K16 acetylation in the protein level **(Fig 6 G)** with increased H4K16 acetylation enriched MMP3 and MMP13 promoter **(Fig 6 H)** and transcriptional upregulation of MMP 3 and 13 **(Fig 6 I)**. These were restored to the level of untreated mice by oral InsP_6_ pretreatment **(Fig 6 G-I)**. While HDAC3 promoter enrichment on the MMP3 and 13 promoter was evident in DSS-treated mice, it was unaltered by oral InsP_6_ pretreatment **(Extended Fig 6 A)**. Its important to mention that oral InsP_6_ also has no effects on IPMK protein expression so works indepdnet of IPMK **(Extended Fig 6B)**.

This finding highlights that disruption of the IPMK-HDAC3 epigenetic axis in a preclinical mouse model of IBD exacerbates intestinal permeability; however, oral InsP_6_ (phytic acid) treatment mitigates the effects of the leaky gut.

### Oral InsP_6_ treatment increased intestinal epithelial cell-specific InsP_6_ level and prevents DSS-induced intestinal injury

To assess the preventative effect of InsP_6_ on intestinal inflammation, wild-type mice were exposed to 2% dextran sodium sulfate (DSS) in drinking water for 7 days, with or without 2% oral InsP_6_ co-treatment **(Fig 7A)**. For the InsP_6_ group, mice were pretreated with oral InsP_6_ for 2 days prior to DSS exposure to enhance the protective effect **(Fig. 7A)**. To induce colitis, mice were given 2% DSS in drinking water continuously for 7 days, followed by sacrifice on the morning of the 8th day **(Fig 7A)**. Each day, mice were also gavaged with 200 microliters of DSS to ensure uniform exposure.

**Figure 7:**
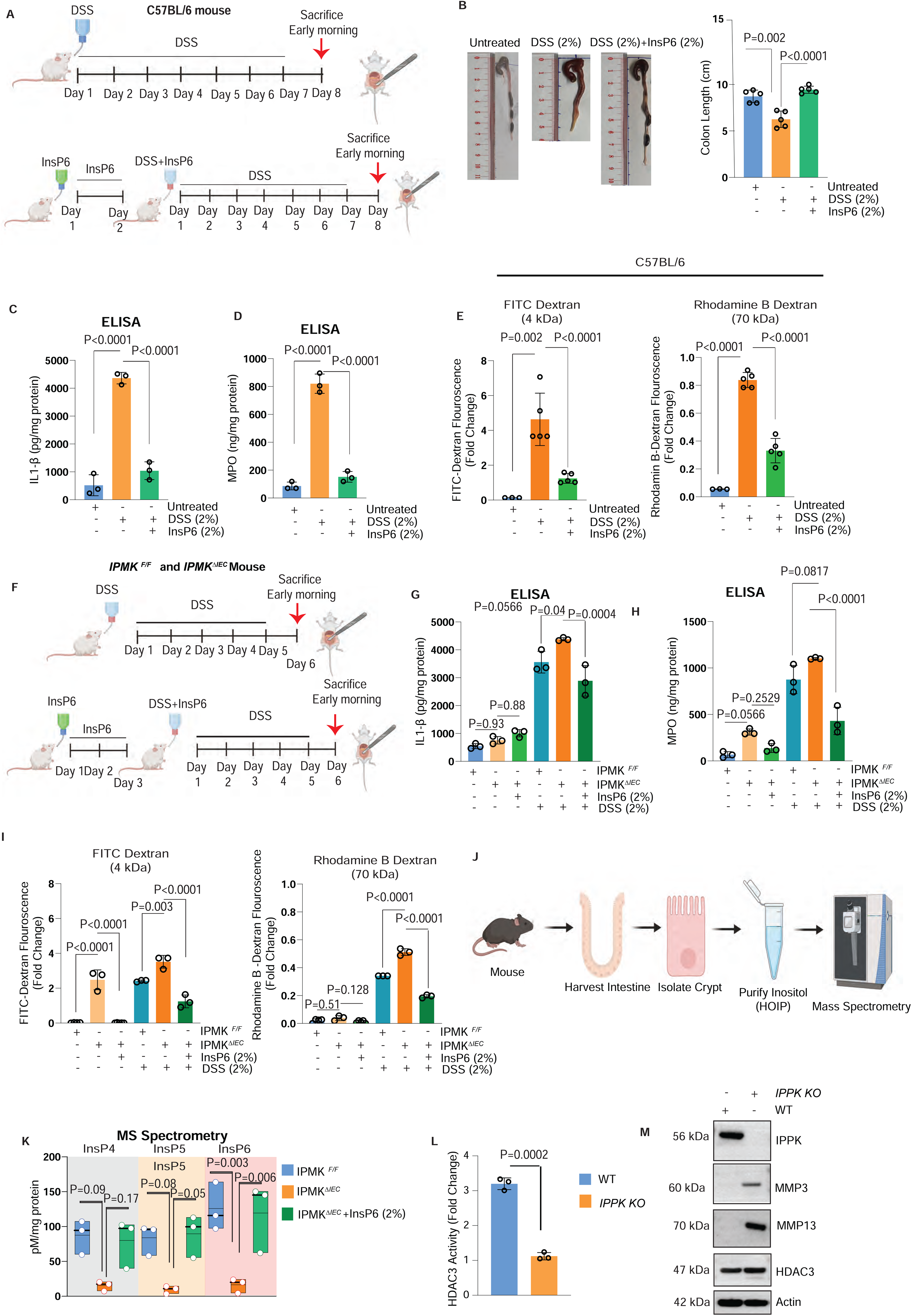
InsP6’s absorption in intestinal tissue prevents DSS-induced inflammation in both wild-type and IPMK^Δ^*^IEC^* mice. **(A)** Schematics of the treatment plan in wild-type mice. **(B)** In wild-type C57BL/6 mice, oral DSS treatment reduces colon size, which InsP6 prevents; n=5. **(C)** ELISA from tissue lysate for IL1β and **(D)** MPO; n=3 for each group. **(E)** Evaluation of intestinal permeability using 4 kDa and 7 kDa dextran. DSS induces intestinal permeability, which oral InsP6 prevents; n=5 in DSS and DSS+InsP6 groups, n=3 untreated. **(F)** Schematics of the treatment plan in IPMK*^F/F^* and IPMK^ΔIEC^ mice. **(G)** ELISA from tissue lysate for IL1β and **(H)** MPO; n=3 for each group, normalized by total protein. **(I)** Evaluation of intestinal permeability using 4 kDa and 7 kDa dextran. DSS induces permeability, which oral InsP6 prevents; n=3 in DSS and DSS+InsP6 groups. **(J)** Schematics of HOIP measurement from intestinal crypts using mass spectrometry. **(K)** Data from mass spectrometry indicates an increase in InsP6 and other HOIPs after oral InsP6 treatment in IPMKΔIEC mice; n=3. **(L)** HDAC3 activity assay in *IPPK* KO cells. **(M)** Western blot of MMPs from *IPPK KO* HEK293T cells.

As expected, mice treated with DSS alone developed severe colitis, characterized by body weight loss starting around 6th day **(Extended Fig. 7A)**, reduced colon length **(Fig. 7B)**, and mucosal erosion and tissue damage observed in H&E-stained colonic sections **(Extended Fig. 7B)**. This inflammatory phenotype was further confirmed by markedly elevated colonic IL-1β (Fig. 7**C**) and myeloperoxidase (MPO) activity (Fig. 7**D**), indicating robust inflammatory cytokine production and neutrophil infiltration, respectively.

Notably, oral co-treatment with InsP_6_ completely prevented these DSS-induced pathologies. Mice receiving InsP_6_ maintained their body weight **(Extended Fig. 7A)** and exhibited no reduction in colon length **(Fig. 7B)** compared to untreated controls. Histological analysis revealed preserved tissue integrity with minimal damage **(Extended Fig. 7B)**, and both IL-1β **(Fig. 7C)** and MPO levels **(Fig. 7D)** remained at baseline, comparable to those of untreated mice. Collectively, these data demonstrate that oral InsP_6_ provides robust protection against the development of DSS-induced colitis.

Lastly, we assessed intestinal permeability. Both 4 kDa and 70 kDa dextran assays showed increased permeability following DSS treatment **(Fig 7E)**, with the greatest increase observed for 4 kDa dextran **(Fig 7E)**. Remarkably, oral InsP_6_ treatment robustly reduced intestinal permeability **(Fig 7E)** and effectively preserved barrier integrity.

We next sought to determine if mice with an intestinal epithelial cell-specific deletion of IPMK (IPMK^ΔIEC^) exhibit heightened susceptibility to DSS-induced inflammation and whether oral InsP_6_ could rescue this phenotype. Three groups of mice were studied: IPMK ^F/F^ (controls), IPMK^ΔIEC^, and IPMK^ΔIEC^ + InsP_6_. All groups were treated with 2% DSS for five days and euthanized on day six **(Fig 7F)**. The IPMK^ΔIEC^ + InsP_6_ group was pre-treated with oral InsP_6_ for three days before DSS co-treatment (Fig 7F), a strategy intended to replenish the depleted endogenous InsP_6_ pools resulting from IPMK loss.

Consistent with a critical role for IPMK in maintaining barrier function, IPMK^ΔIEC^ mice displayed a markedly exacerbated inflammatory response to DSS compared to IPMK ^F/F^ controls, as evidenced by more severe tissue damage in H&E-stained sections **(Extended Fig 7E)** and significantly elevated levels of IL-1β **(Fig 7G)** and MPO **(Fig 7H)**. Furthermore, intestinal permeability to both 4 kDa and 70 kDa dextran was profoundly increased in IPMK^ΔIEC^ mice after DSS treatment **(Fig 7I)**. Remarkably, oral InsP_6_ treatment almost completely abrogated these effects **(Fig 7I)**. The InsP_6_-treated IPMK^ΔIEC^ mice exhibited robust protection, with inflammation markers and intestinal permeability comparable to those of the uninjured IPMK^F/F^ controls. In this experiment the duration of 6 days DSS treatment failed to induce any weight loss or colon size **(Extended Fig 7C,D)**.

While our data established that oral InsP_6_ protects against intestinal inflammation, we sought to determine if this effect stems from direct elevation of intestinal epithelial cell-specific InsP_6_ levels. Mass spectrometry of HOIP metabolites from isolated ileal crypts **(Fig 7J)** revealed that InsP_4_, InsP_5_, and InsP_6_ were markedly reduced in *IPMK*^ΔIEC^ mice relative to IPMK^F/F^ controls, with InsP_6_ levels diminished by approximately sevenfold **(Fig 7K)**. Remarkably, a single 24-hour oral dose of pH-calibrated (7.2) InsP_6_ treated to IPMK^ΔIEC^ mice restored InsP_6_ to levels comparable to controls (Fig 7**K**), while also increasing InsP_4_ and InsP_5_, mirroring observations with cell-permeable InsP_6_ treated to cell lines **(Extended Fig 2C)**. These findings support a model of direct epithelial uptake of orally administered InsP_6_, which is then partially dephosphorylated. Although gut microbial phytases can generate lower-order inositol phosphates and myoinositol for uptake via transporters like SMIT2/SGLT6^34^, the lack of IPMK in IPMK^ΔIEC^ mice would prevent the subsequent re-phosphorylation to InsP_5_ and InsP_6_, thereby implying the existence of a specific epithelial transport mechanism for InsP6 itself.

As we found that incorporation of InsP_6_ at both cellular and tissue levels not only increased InsP_6_ concentration but also elevated InsP_5_ and InsP_4_, we sought to determine which HOIP is physiologically critical for HDAC3 activation and MMP expression. To specifically deplete InsP_6_, we examined HDAC3 activity and MMP expression in *IPPK* knockout cells. IPPK catalyzes the conversion of InsP_5_ to InsP_6_; therefore, the loss of IPPK selectively reduces InsP_6_ without affecting upstream polyphosphates such as InsP_5_ or InsP_4_.

Intriguingly, HDAC3 activity was markedly reduced in *IPPK* knockout cells **(Fig 7L)**, accompanied by elevated MMP levels **(Fig 7M),** however with out influencing HDAC3 expression itself. These findings indicate that InsP_6_ plays a physiologically significant role in HDAC3 activation compared to other HOIPs. In the clinical context, the *IPMK* depletion model better reflects disease conditions such as inflammatory bowel disease (IBD), where IPMK levels are reduced while IPPK remains unchanged **(Fig 5Q,R)**, leading to lowered InsP_6_ concentrations.

## Discussion

Our study provides compelling evidence for the physiological role of InsP_6_ as a crucial activator of HDAC3, with significant implications for epigenetic regulation and intestinal barrier function in IBD. While previous structural studies identified InsP_4_ as a molecular link between HDAC3 and its co-repressors^1^, our findings demonstrate that InsP_6_ is the primary physiological activator of HDAC3, influencing epigenetic changes that regulate MMP gene transcription.

We investigated HDAC3 activation and downstream epigenetic changes in *IPMK*-deleted cell lines and mouse models to determine the physiological role of HOIPs in regulating HDAC3 activation and its impact on the epigenetic axis. We targeted IPMK because it is the rate-limiting enzyme in HOIP biosynthesis^8^, and its deletion resulted in reduced levels of InsP_4_, InsP_5_, and InsP_6_ in cells **(Fig 1B)**. By focusing on IPMK, we aimed to thoroughly evaluate the effects of HOIP depletion on HDAC3-mediated epigenetic regulation.

While *IPMK* deletion markedly diminishes HDAC3’s enzymatic activity **(Fig 1D)**, we discovered that IPMK directly binds to HDAC3 in the chromatin complex **(Fig 1H)**. However, through chemical genetic studies using kinase-dead IPMK and cell-permeable InsP_6_, we revealed that IPMK’s binding to HDAC3 is dispensable for HDAC3’s enzymatic activity **(Fig 2A,B)**. These findings suggest that IPMK’s kinase function, rather than its physical interaction with HDAC3, is critical for maintaining HDAC3’s deacetylase activity. In the physiological context, we hypothesize that IPMK’s localization and association with chromatin-bound HDAC3 may facilitate the generation of HOIPs in specific spatiotemporal regions. This localized production of HOIPs could serve to maintain the efficiency of the HDAC3 regulatory mechanism, ensuring precise control of histone deacetylation in chromatin.

While evaluating the concentration of HOIPs in cells, we made several intriguing discoveries. We found that InsP_4_ concentration was minimal compared to InsP_5_ and InsP_6_ **(Fig 1B).** Notably, *IPMK* deletion primarily diminished InsP_5_ and InsP_6_ levels rather than InsP_4_ **(Fig 1B)**. Consistent with these observations, our *in vitro* rescue experiments demonstrated that both InsP_5_ and InsP_6_ could restore a significant proportion of HDAC3 activity in *IPMK* KO cells, with InsP_6_ exhibiting the highest efficacy **(Fig 2D)**. Furthermore, we observed that cell-permeable InsP_6_ alone was sufficient to restore HDAC3 activity *in vivo* when administered to *IPMK* KO cells **(Fig 2 E)**. These findings collectively underscore the crucial role of InsP_6_ in HDAC3 activation and suggest a direct mechanism of action that is independent of IPMK’s physical presence.

At the mechanistic level, our study revealed that while *IPMK* loss did not disrupt the formation of the HDAC3-corepressor complex **(Fig 3A,B and Extended Fig 3A),** it did impair the DAD (deacetylase activation domain) domain of its corepressor interaction with HDAC3 **(Fig 3D,E),** which is crucial for activating HDAC3’s deacetylase activity however, dispensable for structural integrity.

While HDAC3 recruitment to chromatin is a crucial step in the regulation of gene expression, our study revealed that neither IPMK nor InsP_6_ affects this recruitment process **(Fig 3F, E and Extended Fig 3B)**. Remarkably, we discovered a direct mechanism of HDAC3 activation in a cell-free system. By treating the chromatin fraction from *IPMK* KO cells with InsP_6_, we observed that InsP_6_ facilitates the final step of HDAC3 activation. This activation occurs through the recruitment of the (DAD) domain to chromatin-bound HDAC3, subsequently triggering histone deacetylation **(Fig 3I)**. These findings reveal that InsP_6_ is crucial in activating chromatin-bound HDAC3, serving as a key rate-limiting step. However, InsP_6_ does not contribute to the initial recruitment of HDAC3 to chromatin, highlighting its specific function in enzyme activation rather than chromatin localization.

Building upon our cell model findings that demonstrated a significant role of IPMK and InsP_6_ in HDAC3 activation, we investigated the physiological relevance of the IPMK-HDAC3 epigenetic axis, particularly in the context of IBD, for which IPMK has been identified as a risk gene through GWAS and preclinical studies^13^. Our study revealed that intestinal loss of *IPMK* impaired HDAC3 activity **(Fig 5 I)** in mice, leading to a leaky gut condition **(Fig 5 D)**. At the epigenetic level, disruption of the IPMK-HDAC3 axis increased H4K16 acetylation in MMP promoters **(Fig 4H and Fig 5 J, K)**, activating their transcription **(Fig 4A and Fig 5E)** and resulting in degradation of tight junctions **(Extended Fig 5A)**, thereby exacerbating intestinal permeability **(Fig 5 D)**. Notably, transcriptomic analysis of patients with IBD showed a significant increase in MMPs **(Extended Figure 4 A)**, mirroring observations in intestine-specific *IPMK* knockout mice and suggesting a probable link. Our study further identified that in the mouse model of IBD, the IPMK-HDAC3 epigenetic axis was disrupted due to the loss of IPMK protein expression **(Fig 6 B)**, reflecting the epigenetic effects observed in IPMK^ΔIEC^ mice.

Remarkably, oral administration of InsP_6_ (phytic acid) mitigated the leaky gut effect **(Fig 6A)** by restoring HDAC3 activation **(Fig 6 C)**, reducing H4K16 acetylation **(Fig 6 G)**, and suppressing MMP gene and protein expression **(Fig 6 D - F and I)**, highlighting its potential as an approach in intercepting pre-clinical disease given prior data showing that increased intestinal permeability is associated with later development of Crohn’s Disease. This may also work as a therapeutic approach for IBD, with impaired intestinal permeability shown to be associated with ongoing bowel symptoms and severity of diarrhea in patients with IBD despite having achieved mucosal healing.

Previous studies have indicated that InsP_6_ is absorbed in the gastrointestinal tract, albeit with low bioavailability^25–27,31^. However, studies indicated the therapeutic efficacy of InsP_6_ to treat cancer in mice models^28^. One hypothesis suggests that gut microbiota-derived phytase enzymes dephosphorylate InsP_6_ to lower-order inositol phosphates, which are more readily absorbed^29^. However, our biochemical analysis, corroborated by *Watson* et al.^6^, demonstrates that HDAC3 activity is exclusively influenced by InsP_4_, InsP_5_, or InsP_6_, with InsP_3_ having no effect **(Fig 2 D)**. Notably, oral InsP_6_ administration in IPMK^ΔIEC^ mice, which are incapable of generating InsP_4_, InsP_5_, or InsP_6_, exhibited a rescue effect **(Fig 5 I, J)**. This observation suggests the possibility of direct InsP_6_ absorption in the gastrointestinal tract, although the precise mechanism is yet to be explored.

In conclusion, our study highlights the critical role of the IPMK-HDAC3 epigenetic axis in regulating intestinal barrier integrity and its potential disruption in IBD **(Fig 8)**. We identify InsP_6_ as a key physiological activator of HDAC3 and demonstrate that InsP_6_supplementation can restore HDAC3 activity, reduce histone acetylation, and improve intestinal barrier function in mouse models of IBD **(Fig 8)**.

## Materials and Methods

### Cell culture

Human Colon Tumor 116 (HCT116), Mouse Embryonic Fibroblast (MEF), A549, HEK and CaCo2 cells were cultured in McCoy’s 5A medium, Dulbecco’s Modified Eagle’s Medium (DMEM) or F12/K respectively. DMEM, F12/K media was supplemented with 10% FBS, 2 mM L-glutamine, 100 U/mL penicillin, and 100 mg/mL streptomycin was used for MEF or A549 cells. While McCoy’s 5A medium was supplemented with 10% FBS, 100 U/mL penicillin, and 100 mg/mL streptomycin. *IPMK* wild-type (WT) and knockout (KO) HCT116 and A549 cells were generated using CRISPR/Cas9 technology, which partially deleted exon 5 and the coding region of exon 6. Afterward, clonal populations were isolated, purified, and screened for IPMK expression.

For the generation of *IPMK* KO MEF cells, floxed MEFs were immortalized through transfection with pSG5-Large T (a plasmid provided by William Hahn to Addgene) using the Polyfect transfection reagent (Qiagen) according to the manufacturer’s instructions. After serial passaging five times to select against non-transformed cells, *IPMK* KO cells were produced by transducing floxed cells with an adenovirus carrying the Cre recombinase gene (obtained from the University of Iowa Gene Transfer Vector Core). The virus was combined with Gene Jammer (Stratagene) transfection reagent to improve transduction efficiency. Clonal populations were then purified and screened for IPMK expression.

### Animals

All protocols were approved by the Institutional Animal Care and Use Committee (IACUC, UNLV, IACUC-01204, IACUC-01206), University of Nevada, Las Vegas. Mice were housed according to institutional guidelines, in a controlled environment at a temperature of 22_C ± 1_C, under a 12-h dark-light period and provided with standard chow diet and water *ad libitum*. Male and female *IPMK ^F/F^*, *IPMK*^ΔIEC^ (between eight to 16 weeks) were used. Specifically, intestinal epithelial cells from male mice were used for biochemical and transcriptomic analysis. All mice were maintained in C57BL/6 mixed background.

### Generation of intestine-specific *IPMK* knockout mice

*Ipmk^F/F^* mice were generated as previously described^13^*. Ipmk^F/F^* mice were crossed with C57BL/6J mice carrying Cre expressed under the control of the murine <u>villin</u> promoter as described previously to create intestinal-specific *Ipmk* knockout mice (*Ipmk*^Δ*IEC*^). Homozygous *Ipmk^F/F^* mice were crossed with the *Ipmk*^Δ*IEC*^ mice, which mediate excision of floxed alleles in the intestine. All mice were maintained on a C57BL/6J background and were 10th-generation backcrossed. Validated by MiniMUGA. Background analysis was performed by Transnetyx, Cordova, USA Mice were housed in a 12-hour light/12-hour dark cycle at 22°C and fed standard rodent chow. All research involving mice was approved by the University of Nevada, Las Vegas’s Institutional Animal Care and Use Committee (IACUC).

### Proximity ligation assay and quantitation

PLA was utilized to detect *in situ* protein–protein interactions as previously described. After fixation and permeabilization, cells were blocked with blocking solution supplied by PLA kit before incubation with primary antibodies as in routine immunofluorescence staining. The cells were then processed using a PLA kit (NaveniFlex Cell MR Red Cat no. NAV-NC.MR.100.RED) according to the manufacturer’s instructions. The slides after PLA were further processed for immunofluorescence staining against the DAPI (Cat.8617, Cell Signaling). The slides were then mounted with Prolong Gold antifade Reagent. A Zeiss LSM 750 confocal microscope detected PLA signals as discrete punctate foci and provided the intracellular localization of complexes. ImageJ was used to quantify and represent the number of PLA puncta per nucleus.

### DSS-induced colitis model and oral InsP_6_ treatment

DSS (MP Biomedicals; relative molecular mass 36,000–50,000) was added to drinking water at 3% weight/volume for 24 hr or 2% weight/volume for five or seven days. Phytic acid (InsP_6_) was added to 2% weight/volume (2 gm/100 ml) containing drinking water after pH neutralization with NaHCO for 48 hr after that, the mouse received 2% weight/volume InsP_6_ and 3% DSS for 24 hr or 2% weight/volume for five or seven days..

### Intestinal permeability assay

IPMK*^F/F^* and either IPMK^ΔIEC^ mice were fasted for 4 hours with access to water before being gavaged with 4 kDa-FITC-Dextran and 70 kDa RhodamineB (Sigma-Aldrich, Cat no: #FD4-1G) at a single dose of 0.6mg/gm body weight. Untreated control group received an equal amount of PBS. Fluorescence intensity of plasma samples was measured (excitation 485 nm/emission 535 nm) after 4 hours of gavage. Value was represented as a fold change in respect without FITC Dextran and Rhodamine treated control groups after blank subtraction. All measurements were performed using GraphPad Prism (version 6.0, GraphPad Software, Inc)^22^,.

### Generation of IPMK antibody

Custom anti-IPMK antibody was generated in collaboration with Pro-Sci. The anti-rabbit IPMK antibody had been raised against human IPMK amino acids 295–311 (SKAYSRHRKLYAKKHQS). This sequences are also closely conserved in mice **SK**M**Y**A**RHRK**I**Y**T**KKH**H**S** hence reacts to both mouse and human IPMK.

### Western Blot analysis

Cell lysates were prepared through syringe flush by using a salt-free (NaCl-free) Lysis buffer (50 mM Tris pH 7.5, 50 mM potassium acetate, 5 % v/v glycerol, 0.3 % v/v Triton X-100, one tablet of Roche complete protease inhibitor). We used salt-free (NaCl-free) lysis buffer for all the assays because high salt concentration dissociates higher-order inositol phosphate (HOIPs) from HDAC3, which is essential for their deacetylase activity^25,27^. Samples were centrifuged at 15,000g for 10 min, and the protein concentration of the supernatant was measured. Proteins were resolved by SDS-polyacrylamide gel electrophoresis (NuPAGE Bis-Tris Midi GEL, Life Technologies, Cat #: WG1402BX10) and transferred to Immobilion-P PVDF (Millipore-Sigma, Cat #: IPVH00010) transfer membranes. The membranes were incubated with primary antibody diluted in 3% BSA in Tris-Buffered Saline with Tween 20 (20 mM Tris-HCl, pH 7.4, 150 mM NaCl, and 0.02% Tween 20) overnight incubation at 4°C. Respective antibodies for western blot anti-HDAC3 (Santa Cruz Biotechnology, Cat #: sc-376957), IPMK (Generated in our lab) anti-beta actin (ProteinTech, Cat #: 81115-1-RR), anti-lamin (ProteinTech, Cat #: 12987-1-AP), anti-NCoR1 (Bethyl Laboratories, Cat #: A301-148A), anti-Ncor1 (Cell Signaling Technology, Cat #: 5948), anti-Rcor1 (ProteinTech, Cat #: 26686-1-AP), Sin3A (ProteinTech, Cat #: 14638-1-AP), anti-DNTTIP1 (Bethyl Laboratories, Cat #: A304-048A), anti-H3 (Cell Signaling Technology, Cat #: 12648), anti-H3k8Ac (Cell Signaling Technology, Cat #: 2598), anti-H3k9Ac (Cell Signaling Technology, Cat #: 9649), anti-H3k18Ac (Cell Signaling Technology, Cat #: 13998), anti-H3k27Ac (Abcam, Cat #: ab4729), anti-H3k56Ac (Cell Signaling Technology, Cat #: 4243), anti-H4k8Ac (Cell Signaling Technology, Cat #: 2594), anti-H4k12Ac (Cell Signaling Technology, Cat #: 13944), anti-H4k16Ac (Cell signaling Technology, Cat #: 13534S) and anti-Myc (Invitrogen, Cat #:46-0709). Following primary antibody incubation, the PVDF membrane was washed three times with TRIS-buffered saline/Tween-20, and incubated with HRP-conjugated secondary antibody (ECL, Cat #s: NA934V and NA931V), and the bands visualized by chemiluminescence (Super Signal West Pico, Pierce, Cat #: 34579). The depicted blots are representative replicates selected from at least three experiments. Densitometric analysis was performed using Image J software. Blots for acetylated histones were stripped using Restore Western Blot stripping buffer (Thermo Scientific, Cat # 21059), then reprobed with H3 antibody (Cell Signaling Technology, Cat #: 12648S) at room temperature for 2 h followed by secondary antibody incubation and development.

### HDAC3 (histone deacetylase) activity

To analyze deacetylase activity of specific Class I HDAC members, the HDAC 3 were individually immuno-purified, followed by an activity assay using the following kit: HDAC3 Assay Kit (BPS, Cat#: 10186-628). Since high salt concentration dissociates higher-order inositol phosphates (HOIPs) from HDAC3, which are crucial for their deacetylase activity, WT and IPMK-KO HCT 116 cells were lysed in salt-free (NaCl-free) lysis buffer (50 mM Tris pH 7.5, 50 mM potassium acetate, 5 % v/v glycerol, 0.3 % v/v Triton X-100, one tablet of Roche complete protease inhibitor). Then, the lysate was centrifuged at 15,000 g for 10 min, and the protein concentration of the supernatant was measured. Equal amount, around 20 µg total protein from the lysate was incubated against anti-HDAC3 (Santa Cruz Biotechnology, Cat #: sc-81600) antibodies, respectively, at 4°C for 1 hour followed by capturing antibody with EZview A/G beads (Millipore Sigma, Cat #: E3403). Beads were washed 3x with an ice-cold reaction buffer (supplied with the kit). After a specific substrate reaction, fluorescence was measured using a Spectra max iD5 plate reader (Molecular Devices). IgG was immunoprecipitated from Wild Type HCT 166 cells (WT) and used as a negative control. In each experiment, separate IgG immunoprecipitation was performed, and the data was expressed as a fold change relative to the respective IgG control. Although slight variations in IgG values were observed across experiments, the overall data range differed slightly in each case, but consistently showed a similar trend. All measurements were performed in triplicate, and data was analyzed using GraphPad Prism (version 6.0, GraphPad Software, Inc).

### Intestinal Epithelial cell purification

The ileum (similar length as colon moving proximal from the ileocecal junction) were dissected from mice. After five times washing with ice cold PBS, the tissues were opened longitudinally and cut into small fragments (2–3 cm in length). To prepare single cell suspension after washing three time with PBS, small intestinal fragments were incubated in 30 mM EDTA-PBS on ice for 30 min, during which the tissues were shaken vigorously every 5 min. Isolated crypts were washed once with cold PBS and dissociated using TrypLE Express (Invitrogen Cat# 12604021) for 10 min at room temperature. Cell suspensions were passed through a 70-mm cell strainer and then labeled with a cocktail of fluorescent antibodies specific for APC-CD45 (BioLegend, Cat# 103113), FITC-CD31 (BioLegend, Cat# 102406), PE/Cy7-TER-119 (BioLegend, Cat# 116222), and PE-EpCAM (Thermo Fisher Scientific, Cat# 12-5791-83). Dead cells were excluded using Tail Cell Cycle Solution (Invitrogen, Cat# A10798).Following doublet exclusion and single-cell population isolation through forward and side scatter analysis, the Propidium Iodide^-^ (Ex 535nm: Em 617nm) / APC-CD45^-^ (Ex 651nm: Em 660nm) cell population was gated. Next, FITC-CD31^-^ (Ex 498nm: Em 517nm) / PE-Cy7-TER119^-^ (Ex 565nm: Em 774nm) cells were gated and subtracted from the PI^-^/APC-CD45^-^ population. Remaining cells were gated as PE-EpCAM+ (Ex 565nm: Em 576nm) and isolated as CD45^-^/CD31^-^/TER119^-^/EpCAM^+^ intestinal epithelial cells. CD45^-^/CD31^-^ /TER119^-^/EpCAM^+^ intestinal epithelial cells were sorted by using a SONY SH800S Cell Sorter and collected in a post-sort buffer (RPMI-1640 with 20% FBS).

To isolate crypt from colon, the Colon (from the cecum to the rectum) were dissected out from mice as previously described^33^. After five times washing with ice cold PBS, the tissues were opened longitudinally and cut into small fragments (2–3 cm in length). To isolate crypts after washing three times with cold PBS, small ileum fragments were incubated in 30 mM EDTA-PBS on ice for 30 min, during which the tissues were shaken vigorously every 5 min. Isolated crypts were washed once with cold PBS and dissociated using TrypLE Express (Invitrogen Cat# 12604021) for 10 min at room temperature. Cell suspensions were passed through a 70-µm cell strainer.

### InsP_6_ Rescuing HDAC3 activity

Wild Type and *IPMK*-Knock out (KO) HCT116 cells were lysed in salt-free (NaCl-free) lysis buffer (50 mM Tris pH 7.5, 50 mM potassium acetate, 5 % v/v glycerol, 0.3 % v/v Triton X-100, and one Roche complete protease inhibitor tablet). Samples were centrifuged at 15,000 g for 10 min, and the protein concentration of the supernatant was measured. Equal amount, 20 µg total protein from the lysate was incubated against anti-HDAC3 antibodies respectively at 4°C for 1 hr, followed by capturing antibodies with EZview A/G beads (Millipore Sigma, Cat #: E3403). Beads were washed 3x with an ice-cold reaction buffer (supplied with the kit). Following washing, beads were resuspended in reaction buffer then beads were incubated with 1nM, 10nM, 100nM, 500nM, 1µM, 5µM and 10µM concentrations of InsP_6_ (Sigma Aldrich Cat #: P8810-100G) for 1 hr at room temperature. After incubation, beads were washed 3x with a reaction buffer. HDAC3 activity was measured by the addition of a fluorogenic substrate provided by the HDAC Assay Kit (BPS, Cat #: 10186-628) and read using a Spectramax iD5 plate reader (Molecular Devices). IgG was immunoprecipitated from IPMK Wild Type HCT 166 cells (WT) and used as a negative control. In each experiment, separate IgG immunoprecipitation was performed, and the data was expressed as a fold change relative to the respective IgG control. Although slight variations in IgG values were observed across experiments, the overall data range differed slightly in each case, but consistently showed a similar trend. All measurements were performed in triplicate data and analyzed using GraphPad Prism (version 6.0, GraphPad Software, Inc)^15,13,16^.

### Analysis of rescuing HDAC3 activity using PIP3, InsP_3_, Ins (1,3,4,5) P_4_, Ins(1,4,5,6)P_4_, InsP_5_ and InsP_6_

Wild type and *IPMK*-Knock out (KO) HCT116 cells were lysed in salt-free (NaCl-free) lysis buffer (50 mM Tris pH 7.5, 50 mM potassium acetate, 5 % v/v glycerol, 0.3 % v/v Triton X-100, and one Roche complete protease inhibitor tablet). Samples were centrifuged at 15,000 g for 10 min, and the protein concentration of the supernatant was measured. Equal amount, 20 µg total protein from the lysate was incubated against anti-HDAC3 antibodies respectively at 4°C for 1 hr, followed by capturing antibodies with EZview A/G beads (Millipore Sigma, Cat #: E3403). Beads were washed 3x with an ice-cold reaction buffer (supplied with the kit). Following washing, beads were resuspended in reaction buffer then beads were incubated with either 10µM concentration of PIP3, InsP_3_, Ins(1,3,4,5)P_4_ or 1µM Ins(1,4,5,6)P_4_, InsP_5_ and InsP_6_ (Sigma Aldrich Cat #: P8810-100G) for 1 hr at room temperature. After incubation, beads were washed 3x with a reaction buffer. HDAC3 activity was measured by the addition of a fluorogenic substrate provided by the HDAC3 Assay Kit (BPS, Cat #: 10186-628) and read using a Spectramax iD5 plate reader (Molecular Devices). IgG was immunoprecipitated from Wild Type HCT 166 cells (WT) and used as a negative control. In each experiment, separate IgG immunoprecipitation was performed, and the data was expressed as a fold change relative to the respective IgG control. Although slight variations in IgG values were observed across experiments, the overall data range differed slightly in each case, but consistently showed a similar trend. All measurements were performed in triplicate data and analyzed using GraphPad Prism (version 6.0, GraphPad Software, Inc)^15,13,16^.

### Plasmids and Recombinant Proteins

pMX-myc, pMX-*IPMK-WT-*myc and pMX-*IPMK-KSA*-myc plasmids transiently transfected into *IPMK*-KO HCT 116 cells using Lipofectamine 3000 (Cat no. L3000001, Thermo Scientific) according to the manufacturer protocol. Recombinant myc-IPMK was purchased from Origene, (Cat #: TP309343) while recombinant GST-HDAC3 (Cat #: H85-30G) and GST (Cat #: G52-30U) were purchased from Signal Chem.

### Cell fractionation and isolation of chromatin fraction

Approximately 7×106 cells were isolated and washed three times with chilled PBS. After that cells were lysed with ice cold cell lysis buffer (50 mM HEPES-KOH, 140 mM NaCl,1 mM EDTA, 10% glycerol, 0.5% NP-40, 0.25% Triton X-100, 1 mM DTT Supplemented with 1x protease inhibitor cocktail). After centrifugation at 1100 g for 5 min supernatant was separated as a cytoplasmic fraction. Pellets were the crude nuclei. After washing three time with cell lysis buffer, nuclei was incubated 10 min in ice with nuclear lysis buffer (10 mM Tris-HCl, 200 mM NaCl, 1 mM EDTA, 0.5 mM EGTA, Supplemented with 1x protease inhibitor cocktail). Gently resuspended 10 times. Then centrifuge at 1100 g for 10 min. Supernatant was separated as a nuclear fraction and pellet was the crude chromatin. After washing three times with a nuclear lysis buffer, crude unsheared chromatin was used for further experiment.

### Immunoprecipitation

To analyze endogenous binding, HCT116 cells were lysed in salt-free lysis buffer (50 mM Tris pH 7.5, 50 mM potassium acetate, 5 % v/v glycerol, 0.3 % v/v Triton X-100, and one Roche complete protease inhibitor tablet). For the endogenous immunoprecipitation (IP), respective antibodies against HDAC3 were used, followed by western blot of the binding partners. In brief, the IP was performed from 500 μug of protein lysate. Protein lysates were incubated for 2 hr at 4°C with respective antibodies, then the antibody-protein complex was captured with EZview A/G beads (Millipore Sigma, Cat #: E3403). Beads were pelleted and washed with lysis buffer 3x, followed by elution in sample buffer. Immunoprecipitated samples were resolved on a NuPAGE Bis-Tris gel, followed by western blotting ^18^

### ChIP-qPCR

Approximately 7×106 cells were fixed with 1% final volume from fresh 16% stock formaldehyde (Sigma-Aldrich, Cat#: F8775) at room temperature for 10 min followed by ChIP using ChIP assay Kit from iDEAL Diagenode ( Cat no.C01010051). Cells were then harvested and lysed in 500 mL of ChIP lysis Buffer (50 mM Tris-HCl pH 8.0, 5 mM EDTA, 150 mM NaCl, 0.5% Triton X-100, 0.5% SDS, 0.5% NP-40, 1 mM sodium butyrate) containing protease inhibitor cocktail. The lysates were subjected to sonication to shear DNA to a length of approximately between 150 and 900 bp. The lysate was then diluted in 0.75 mL of ChIP dilution buffer and incubated with control IgG (Cell Signaling Technology, Cat#: 2729S) or primary antibody H3K9 ac (Cells signalling Technology, Cat#9649T), H3K27 ac (Abcam cat# 4729), H4k16ac (Cells signalling Technology, Cat#13534), HDAC3 ( Santa Cruz Biotechnology, Cat# 39717) and HDAC1 ( Santa Cruz Biotechnology, Cat# 81598X) 4C overnight. Then the lysate was incubated with Protein G magnetic bead (provided in kit) for 1 h at 4C. The beads were washed sequentially with a wash buffer provided in the kit. The immune complexes were eluted with 75 mL of elution buffer (1% SDS, 0.1 M NaHCO3) twice at 65C for 30 min. After elution, the cross-link was reversed by adding NaCl and incubated together with Proteinase K provided with the kit overnight at 65C. ChIP DNA was purified using the ChIP DNA purification kit Provided with the kit. The purified DNA was analyzed on a StepOnePlus using the power SYBR Green Master Mix. The results are presented as a fold change in respect with IgG after calculating the percentage of input. qPCR analyses were done in triplicate. We used MMP3 5′-AAAATAGAGTAGCAGAGGCAGGTA −3′ and 5′-AGAGTGGTGGCAGTGATGTGAA −3′, and MMP 13 Fwd 5′ CAACCATGGGGCTCAATCCT3′ Rev 5′ CTTACGTGGCGACTTTTTCTTTTC3′ Primer for human and MMP3 5′-TGTGGTCACTGATAGTGTGGACTGT −3′ and 5′-CCAGAAGCCCAAGCTGAATACTGT −3′, and MMP 13 Fwd 5′ CTGCCACAAACCACACTTAGG3′ Rev 5′ AGTCACCACTTTGGGGTGTG3′ Primer for mouse ^18^

### mRNA expression by qPCR

After extracting the total RNA using RNeasy Mini Kit (Qiagen), and checking its integrity by nanodrop, the cDNA was synthesized from 1 mg of purified total RNA using High capacity cDNA Reverse Transcription Kit (Applied Biosystem). Expression of mouse and human MMP1, MMP3 and MMP13, 18S for human and mouse GAPDH was detected using suitably designed Taqman primers (Invitrogen). The experiments were performed (real-time PCR Systems StepOne plus, Applied Biosystems) in triplicate. Data were quantified for the above genes using the comparative Ct method, as described in the Assays-on-Demand Users Manual (Applied Biosystems)^18^

### *In vitro* Binding assay

Equal amount of recombinant myc-IPMK (Origene, Cat #: TP309343) was co-incubated with either GST-HDAC3 (Signal Chem, Cat #: H85-30G) or GST (Signal Chem, Cat #: G52-30U) in lysis buffer and the complex was maintained for 30 min at 4°C. After the addition of GST beads, incubation continued for an additional 45 min and washed 3x with an ice cold lysis buffer. SDS sample buffer was added, and binding was confirmed by western blotting of anti-Myc antibodies^18^.

### Transwell assay

Transwell filter inserts (pore size 1.0 μm, 12 mm diameter, polyester membrane, Corning, New York, USA) were coated with fibronectin (10 μg/ml; Sigma) used as insert for 24 well plate. For time dependent permeability assay CaCo2 cells were seeded (0.25 × 10^6^ cells/well) and grown on transwell filters for 48 hours until reaching confluency. FITC-dextran 4 kDa-FITC-Dextran (Sigma-Aldrich, Cat no: #FD4-1G) was added to the top well at 25 mg/ml. at 37°C, and fluorescence was measured after 10 min by using a Spectramax iD5 plate reader (Molecular Devices, (excitation 485 nm/emission 535 nm)^47^.

To analyze crucially of IPMK or InsP_6_in cell permeability, CaCo2 cells were seeded (0.25 × 10^6^ cells/well) and grown on transwell filters for 48 hours until reaching confluency. Soup from WT and *IPMK*-KO HCT116 cells and confluent monolayers of WT and *IPMK*-KO HCT 116 cells were treated with similar amounts of PBS (as a negative control), Ilomastat (10µM) or cell permeable InsP_6_(50µM) or MMP Activator p-Aminophenylmercuric acetate (5µM) overnight. Soup from those HCT 116 cell cultures was transferred to mono layer of CaCo2 cells in the transwell plate. Incubate for four hours. FITC-dextran 4 kDa-FITC-Dextran (Sigma-Aldrich, Cat no: #FD4-1G) was added to the top well at 25 mg/ml for 10 min at 37°C, and fluorescence was measured using a Spectramax iD5 plate reader (Molecular Devices, (excitation 485 nm/emission 535 nm).

### RGFP966 treatment

One million cells WT and *HDAC3* or *IPMK* knockout HCT 116 cells were plated and let them grow overnight. Next day cells were treated with HDAC3 specific inhibitor RGFP966 at a dose of 10µM for 24 hr.

### CP-InsP_6_ treatment

Cell permeable InsP_6_ was created and characterized in Prof. Dr. Henning J. Jessen Lab, University of Freiburg, Germany^17^. After synthesis cell permeable InsP_6_ was primarily dissolved in DMSO to prepare a 10 mM primary stock. One million cells *IPMK* KO HCT 116 cells were plated and let them grow overnight. Next day cells were treated with CP-InsP_6_ at a dose of 50µM for 24 hr in media.

### Intracellular inositol content analysis using HPLC

*IPMK*-WT and *KO* HCT116 cells were plated at a density of 250,000 cells per 60 mm plate, then labeled with 60 mCi (1 Ci = 37 GBq) [3H] myo-inositol (PerkinElmer) in conventional cell culture media for three days. To extract soluble inositol phosphates, cell pellets were suspended in 300 mL of ice-cold 0.6 M perchloric acid buffer (0.1 mg/mL IP6, 2 mM EDTA) and incubated on ice for 1 min. Ninety mL of 1 M potassium carbonate with 5 mM EDTA were added and incubated on ice for 1 h. Extracts were centrifuged at 12,000 rpm for 15 min. The supernatant was collected and analyzed by HPLC using a Partisphere SAX column (Whatman Inc.). The column was eluted with a gradient generated by mixing Buffer A (1 mM EDTA) and Buffer B (Buffer A plus 1.3 M (NH4)2HPO4, pH 3.8 with H3PO4). The 1 mL fractions were collected and counted using 5 mL of Ultima-Flo AP mixture (PerkinElmer).

### Unlabelled Inositol phosphates purification from cells and tissue

Inositol phosphates (HOIP) purification was performed as previously described^35^. Briefly, for cell culture HCT116 cells total of around 30-40 million) was used and the total inositol content was normalized by the total cell count. The data is represented as picomole (pM) per 1 million cells. For the mouse intestinal crypts from the Ileum they were washed in cold PBS, and a fraction was set aside for normalization using total protein. The samples were resuspended with cold 1 M perchloric acid and incubated on ice. Titanium dioxide beads were prepared as described (4 mg/sample). Samples were centrifuged and the supernatant was transferred to Eppendorfs containing 4 mg of titanium dioxide beads and rotated at 4°C for 20 minutes. Samples were centrifuged, and the beads were washed with cold 1 M perchloric acid three times. Inositol phosphates were eluted twice by resuspending in ammonium hydroxide and rotating at 4 C for 10 minutes. Purified inositol phosphates were dried on low heat using a centrifugal evaporator.

### Inositol phosphates quantification using mass spectrometry

The analysis was performed on a CE-QQQ system (Agilent 7100 CE-with Agilent 6495C Triple Quadrupole and Agilent Jet Stream electrospray ionization source, adopting an Agilent CE-ESI-MS interface). An isocratic Agilent 1200 LC pump was used to deliver the sheath liquid (50% isopropanol in water) with a final splitting flow speed of 10 µl/min via a splitter. All separation was performed with a bare fused silica capillary with a length of 100 cm (50 µm internal diameter and 365 µm outer diameter). 40 mM ammonium acetate titrated with ammonium hydroxide to pH 9.0 was used as background electrolyte (BGE). Between runs of each sample, the capillary was flushed with BGE for 400s. Samples were injected by applying 100 mbar pressure for 15s (30 nL). The MS source parameters were as follows: gas temperature was 150°C, gas flow was 11 L/min, nebulizer pressure was 8 psi, sheath gas temperature was 175°C and with a flow at 8 L/min, the capillary voltage was −2000V, the nozzle voltage was 2000V. Negative high-pressure RF and negative low-pressure RF were 70 V and 40 V, respectively. Set multiple reaction monitoring (MRM) as shown in the table below.

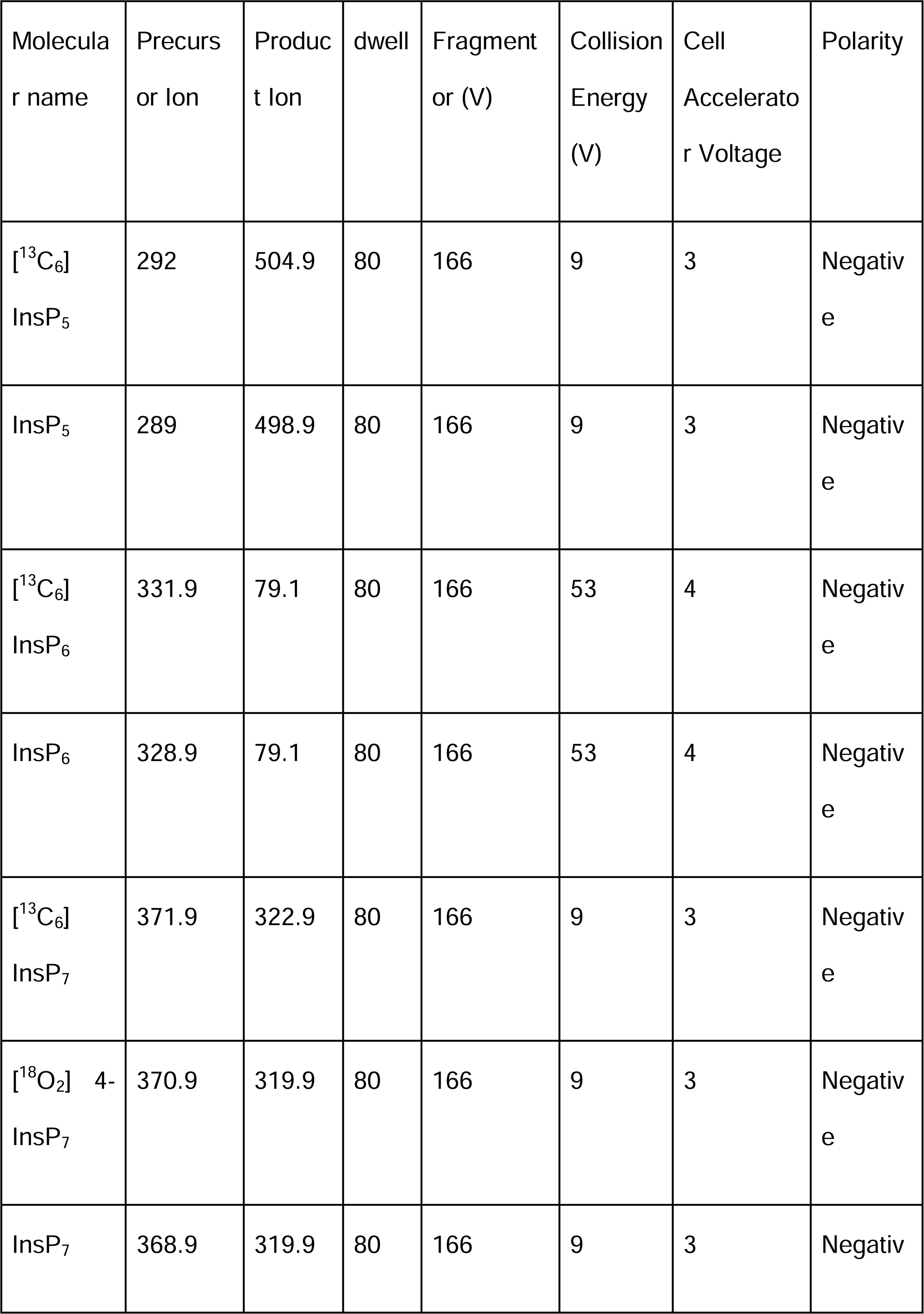
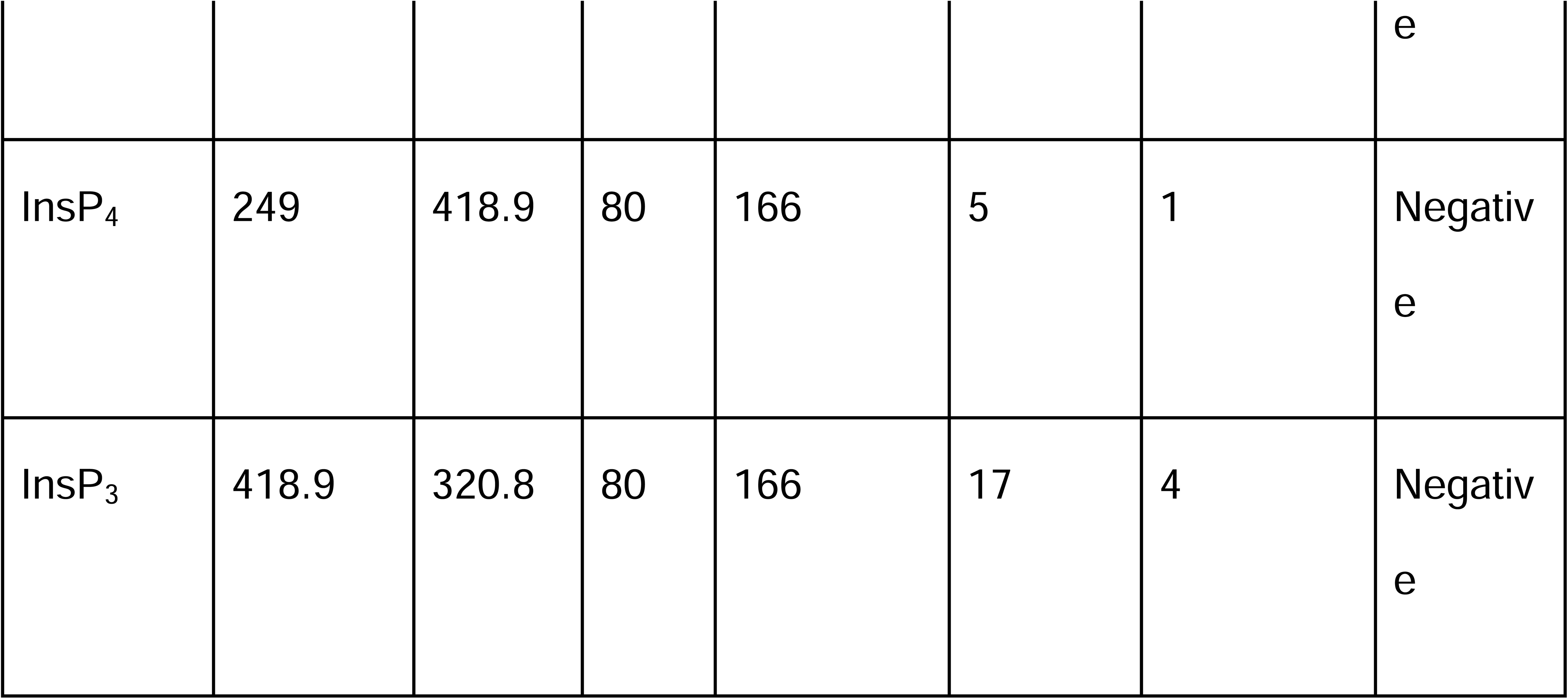

Internal standard (IS) stock solution of were spiked to samples for the assignment and quantification of InsPs and PP-InsPs. Quantification of 1,5-InsP_8_, 5-InsP_7_, 1-InsP_7_, and InsP_6_ was performed with known amounts of corresponding heavy isotopic references spiked into the samples. Quantification of 4/6-InsP_7_ was performed with 13C 5-InsP_7_ and Quantification of InsP_3-4_ of which no isotopic standards are available was performed with spiked 13C InsP_6_. After spiking, 20 μM [^13^C_6_] 2-OH InsP_5_, 20 μM [^13^C_6_] InsP_6_, 4 μM [^13^C_6_] 5-InsP_7_, 2 μM [^13^C_6_] 1-InsP_7_ and 1 μM [^13^C_6_] 1,5-InsP_8_ were the final concentration inside samples. All 13C inositol references were kindly provided by Dorothea Fiedler^36^.

### Histology and Immunohistochemistry

10% Phosphate-buffered-paraformaldehyde fixed intestine and colon tissues was subsequently opened lengthwise and washed in cold PBS, then cut into approximately 3-4 mm fragments for sectioning and paraffinization. After that tissue was dehydrated through 70%, 90% 100% and xylene alcohol gradation and subjected to block preparation in low melting point (65°C) paraffin wax. six micron tissue sections were cut and spreaded over Mayer’s albumin coated slides. Tissue slides were deparaffinized and stained with hematoxylin and eosin following standard protocol for histological evaluation under light microscope.

Mice were euthanized by CO_2_. Immediately following euthanasia, the intestine and colon was removed and flushed with cold PBS. The intestine was subsequently opened lengthwise and washed in cold PBS, then cut into approximately 3-4 mm fragments for sectioning and paraffinization. Samples were deparaffinized with Histo-Clear (Thermo Fisher) and rehydrated in successive washes of 100%, 90%, and 75% ethanol followed by deionized (DI) water. Sections were unmasked via heat-induced epitope retrieval (HIER) using citrate buffer, followed by blocking with 3% goat serum for 1 hour. Primary antibodies were diluted per manufacturer’s recommendation in 1% Triton-X containing blocking solution, incubating overnight at 4°C in a humidified chamber. The following day, samples were washed with TBST, followed by incubation with fluorescent-tagged secondary antibodies for 60 minutes at 37°C. After five time washing with TBST, samples were stained with 4′, 6-diamidino-2-phenylindole (DAPI) diluted in PBS (1 μg/mL). Slides were imaged using an Axiovert Zeiss epifluorescence microscope and LSM800 confocal microscope. Subsequent histogram analysis in ImageJ produced signal intensities from 0-255, with 0 = darkest intensity (positive staining) and 255 representing a white, unstained signal. A threshold of 0-230 was applied to eliminate background staining.

### RNA isolation and library preparation

Total RNA form was isolated by using RNAeasy Mini Kit (Qiagen Cat no.74104) by following manufacturer’s kit protocol. For RNA-seq, libraries were prepared from total RNA by using True Seq RNA Library Prep Kit V2 (Illumina).

### RNA seq analysis

FASTQ files were retrieved from sequencing. All pre-alignment and post-alignment QA/QC and data analysis were performed in Partek Flow (v.12.0.0). Reads were aligned to the whole genome using hg19 assembly with STAR (v.2.7.8a). Aligned reads were quantified to the hg19 annotation model using HTSeq (v.0.11.0). Differential analysis was performed for *IPMK* knockout versus wild-type, *HDAC3* knockout versus wild-type, and IPMK^ΔIEC^ versus IPMK*^F/F^* using DESeq2 (v.3.5)31.

### Pathway analysis using Metascape

Pathway analysis was performed on differentially expressed genes (FDR <= 0.05 and log2FoldChange >= 2) for HCT116 IPMK knockout versus wild-type and Vilcre versus FF using Metascape (v.3.5) with default settings. The genes were queried against GO Biological Pathways, KEGG Pathway, Reactome Gene Sets, Canonical Pathways, WikiPathways, and Panther pathways. Terms with a p-value < 0.01, a minimum count of 3, and an enrichment factor > 1.5 were considered. All genes were used as a background gene list.

### GSEA pathway analysis

Gene set enrichment analysis (GSEA) was performed using the R package cluster Profiler^37^. We used all the genes after DESeq2 normalization to generate a ranked gene list sorted by Wald’s test statistic, then queried relevant pathways containing MMP genes from the C2:CP:Reactome subcollection of the Molecular Signature Database (MSigDB)^38^.

### ELISA analysis of ILβ and MPO

ELM-IL1b-1, ELM-MPO-1). Briefly, tissue lysates were incubated in wells precoated with the relevant antibody, and detection was carried out by measuring absorbance at 450nm. Final optical density values were blank-subtracted and normalized to total protein content.

### Statistical analysis

All plots and statistical analyses were performed with Prism 9 (GraphPad) software. Statistical significance was determined by either Student’s t-test (two-tailed) for two groups or 1-way ANOVA for multiple groups with similar samples. Error bars represent the standard deviation of the mean and indicate replicates or the number of animals employed. Results were representative of at least three independent experiments (n). Graphs were generated by using either Prism 9 (GraphPad) software or R software.

## Reporting summary

Further information on research design is available in the Nature Portfolio Reporting Summary linked to this article.

## Data availability

Once the paper published the raw data will be included as supplemental figures and genomic data will be uploaded in GEO platform.

## Acknowledgements

This work was supported by NIH R16GM154726 and 5P20GM121325 COBRE grant and University of Nevada, Las Vegas start-up funds to Prasun Guha. This work was supported by UK Research and Innovation (UKRI)’s Medical Research Council (MRC) grant MR/T028904/1. We would like to thank NIPM’s Genomic Core for assisting with instruments and experiments. The specimen collection and phenotyping was made possible by the Washington University (WU) DDRCC (NIDDK P30 DK052574). Parakkal Deepak was supported by the Helmsley Charitable Trust, the IBD Plexus of the Crohn’s and Colitis Foundation, and the Leo & Carean Goss Crohn’s Disease Research Fund. We would like to thank Marc C Jhonson, from University of Missouri-School of Medicine for generously providing *IPPK* Knock out cells. We would like to thank Mark Donowitz from Johns Hopkins Medicine, Subrata H Mishra from NIST and Brian Hedlund from UNLV for reading the paper and giving suggestions.

## Author Contributions

PG conceived the study. PG and SC designed the experiments. SC performed most experiments. LVP validated major biochemical experiments. SC performed wet-lab experiments related to NGS studies. ZS and LVP performed most of the NGS data analysis. KR and HJJ generated the cell-permeable InsP_6_. XBS and AS analyzed intercellular inositol content. NT and SC performed animal and microscopy related experiments. QL performed NGS data validation. SJR maintained animal colonies and isolated animal samples for genotyping. SP performed mathematical analysis of transwell assay. SC and ZS designed the Figures. PG and SC wrote the manuscript. SC takes responsibility of all wet lab data.

**Extended Figure 1: (A)** Western blot analysis of the lysates showed complete loss of IPMK protein in three CRISPR Knock out cell lines (HCT116, MEF and A5490 as compared to lysate isolated from respective Wild type (WT) HCT116 cells. Actin was used as a loading control. Results were representative of three individual experiments (n=3). **(B)** HDAC3 were immunoprecipitated from 5µM, 20µM, 50µM, 100µM and 500µM Wild Type (WT) HCT 116 cell lysate followed by HDAC3 activity assay. Data was represented as a fold change compared to IgG. Results are representative of minimum three individual experiments. (n=3) **(C)** Western blot analysis of the lysates isolated from WT and *IPMK* Knock out (KO) HCT 116 cells showed HDAC3 and its corepressor protein NCoR1/2 levels were comparable in all genotypes. Actin was used as a loading control. Results were representative of three individual experiments (n=3). **(D)** Western blot analysis of the lysates showed complete loss of HDAC3 protein in CRISPR Knock out cell lines HCT116, as compared to lysate isolated from respective Wild type (WT) HCT116 cells. Actin was used as a loading control. Results were representative of three individual experiments (n=3). **(E,F)** Western blot analysis demonstrated that histone acetylation increased in *IPMK-* Knock out -HCT 116 cells (KO) compared to the Wild Type HCT 116 cells (WT). *IPMK* deletion specifically increased acetylated H3K9, and 27 and H4k16. Marked by *, denoted high alteration level. Data represents three experimental replicates (n=3). Followed by densitometric analysis. **(G)** After incubation with recombinant HDAC3-GST, and IPMK-myc the sample was immunoprecipited with myc bead and western blotted against anti-GST antibody. Result showed recombinant IPMK binds with HDAC3. GST protein was used as control. Results was representative of three experimental replicates (n=3).

**Extended Figure 2: (A)** Next, efficacy of 0.01µM InsP_4_ and InsP_5_ analyzed compared to InsP_6_ in activating HDAC3 activity *in vitro*. Data was represented as a fold change compared to respective IgG. Results were representative of three independent experiments (n=3). **(B)** 50 µM cell permeable InsP_6_ for 24 hr failed to increase HDAC3 activity when treated to WT cells. Data was represented as a fold change compared to IgG. Results are representative of three individual experiments (n=3). **(C)**Mass spectrometric analysis of HOIP level after IPMK KO HCT116 cells treated with CP-InsP_6_ for 24 hours.(n=3)

**Extended Figure 3 I: (A)** Immunoprecipitation of HDAC3 followed by western blot of various corepressor proteins (NCoR1/2) indicates that the interaction of HDAC3 and NCoR1/2 in *IPMK* KO HCT 116 cells was comparable to WT HCT 116 cells. The result was representative of three individual experiments. (n=3). **(B)** The interaction of HDAC3 and H3/H4 in IPMK KO HCT 116 cells was comparable to WT HCT 116 cells. The result was representative of three individual experiments. (n=3). **(C)** Immunoprecipitation of NCoR2/1 from cell free chromatin followed by western blot of HDAC3, GPS2, TBL1 indicates that the interaction of HDAC3, TBL1, GPS2 with NCoR1/2 in *IPMK* KO HCT 116 cells was comparable to WT HCT 116 cells. Immunoblotting against anti RNA POL 2 (chromatin) and E Cadherin (cytoplasmic and cell membrane) antibody used as cell fractionation control. Data represents three experimental replicates (n=3).

**Extended Figure 3 II: (A)** Immunoprecipitation of HDAC3 followed by western blot of IPPK indicates that the interaction of HDAC3 and IPPK in *IPMK* KO HCT 116 cells was comparable to WT HCT 116 cells. The result was representative of three individual experiments. (n=3). **(B)** Proximity ligation assay using anti-HDAC3 and anti-IPPK antibodies showed RED PLA puncta in the nucleus of WT HCT 116 cells. Scale bar 5μM (n=35).

**Extended Figure 4 I: (A)** Volcano plot (Log 10[*FDR*] vs. Log 2[Fold change]) displays differentially expressed genes in Crohn’s disease patients from the Inflammatory Bowel Disease Transcriptome and Metatranscriptome Meta-Analysis (IBD TaMMA) database as compared with healthy individuals. Red dots represent gene expression upregulated in Crohn’s patient ileum, while blue dots represent down regulated gene expression. The Y-axis denotes the –Log10 FDR Value, while the X-axis denotes the Log2 fold change value. The result represented an expression of various MMP genes high in Crohn’s disease patient’s ileum. **(B)** *IPMK*-Knockout (KO) HCT116 cells transiently overexpressed with *myc-control*, *wt-IPMK*myc, and *KD-IPMK-*myc plasmid. Western blot analysis demonstrated that *myc-control* and *KD-IPMK-*myc *were* unable to rescue MMP1, MMP10 or MMP13 expression while wt-IPMK-myc was rescued. Actin, was used as a loading control respectively. Results were representative of three independent experiments (n=3). **(C)** qPCR analysis of mRNA isolated from WT and *HDAC3* KO HCT 116 (HDAC3 KO) cells indicated increased MMP1, MMP3 and MMP13 gene transcription. 18s used as loading control. Results represented three experimental replicates (n=3). **(D)** ChIP-qPCR analysis using anti-HDAC3 and anti-HDAC1 antibodies showed MMP13 promoter enrichment was exclusively on HDAC3 and comparable in WT and *IPMK* KO cells. Data represents three experimental replicates (n=3). **(E)** qPCR analysis of mRNA isolated from WT and *IPMK* KO HCT 116 (KO) cells indicated increased MMP3 transcription. 18s used as loading control. Results represented three experimental replicates (n=3). **(F)** Western Blot analysis with anti-MMP3 antibody showed profound increase in MMP3, expression at protein level after *IPMK* deletion in HCT 116 cells (KO) as compared to WT and **(G)** was comparable to *HDAC3* KO HCT 116 cells (HDAC3 KO). Actin used as loading control. Data represents three experimental replicates (n=3). **(H)** Western blot analysis revealed that the deletion of *IPMK or HDAC3* led to profound increased MMP3 expression as compared to WT HCT116 cells. An 8-hour treatment with the HDAC3-specific inhibitor RGFP966 at a dose of 10 µM in HCT116 WT, IPMK KO, and HDAC3 KO cells showed no further increase in MMP3 expression compared to the untreated *IPMK* KO and *HDAC3* KO HCT116 cells. Actin used as loading control. Data represents three experimental replicates (n=3). **(I)** Western blot of MMP3 from IPMK WT and KO cells. 24 hr of 50 µM CP-InsP_6_ treatment to KO cells successfully rescued MMP3 expression while co treatment with 10 µM RGFP966 abrogated InsP_6_’s rescue effect. Actin used as loading control. Data represents three experimental replicates (n=3).

**Extended Figure 4 II: (A,B)** Volcano plot (Log 10[*FDR*] vs. Log 2[Fold change]) displays differentially expressed genes in Crohn’s disease and ulcerative colitis patients from the Inflammatory Bowel Disease Transcriptome and Metatranscriptome Meta-Analysis (IBD TaMMA) database as compared with healthy individuals. Red dots represent gene expression upregulated in patient tissue, while blue dots represent downregulated gene expression. The Y-axis denotes the –Log10 FDR Value, while the X-axis denotes the Log2 fold change value. The result represented increased expression of various MMP genes.

**Extended Figure 5 I: (A)** Monolayer of CaCo2 cells were treated with conditioned media from WT and *IPMK* KO HCT 116 cells for 8 hr. After treatment CaCo2 cells were lysed and western blotted against anti-ZO-1 and anti-Occludin antibodies, actin was used as loding control, followed by densitometric analysis. Data represents three experimental replicates (n=3). **(B)** Pictorial representation of FACS based intestinal epithelial cell purification procedure. **(C)** After isolation, intestinal cells were flow purified, followed by staining with APC-CD45 (Ex 651nm: Em 660nm), FITC-CD31(Ex 498nm: Em 517nm), PE/Cy7-TER-119 (Ex 565nm: Em 774nm), and PE-EpCAM (Ex 565nm: Em 576nm). Dead cells were excluded using the Tail Cell Cycle Solution, propidium iodide (PI) (Ex 535nm: Em 617nm). CD45^-^/CD31^-^/TER-119^-^ /EpCAM^+^ intestinal epithelial cells were sorted using a SONY SH800S Cell Sorter. **(D)** CD45^-^/CD31^-^/TER-119^-^/EpCam^+^ intestinal epithelial cells were isolated from IPMK^Δ^*^IEC^* and IPMK *^F/F^* mice through FACS purification showing loss of IPMK protein in IPMK^Δ^*^IEC^ mice.* Actin was used as a loading control. **(E)** Western blot analysis showed the purity of EpCAM ^+^ intestinal epithelial cell population. Lysate was isolated from unsorted and sorted intestinal cells, followed by western blot against anti-CD31, anti-CD45, anti-TER-119, and anti-EpCam. Actin was used as a loading control. Analysis showed that the sorted cell population was devoid of CD31, CD45, and TER-119 cell population but positive for EpCam. Results were representative of three individual experiments (n=3). **(F)** Western blot from intestinal crypts (ileum) indicated elevated H4k16 acetylation in IPMK^ΔIEC^ mice, which was reduced by pretreatment of 2% oral InsP_6_. Western blot analysis further showed IPPK protein levels were comparable in all untreated and untreated genotypes. Lamin was used as a loading control. Results were representative of three individual experiments (n=3). (G) Western blot analysis of colonic crypts revealed elevated levels of MMP3, MMP13, and acetylated H4K16 in IPMK^Δ^*^IEC^* mice, which were rescued following pretreatment with 2% oral InsP6. IPMK expression was diminished, while IPPK levels remained unaltered. Actin served as a loading control. Data are of three independent experiments (n = 3). **(H)** ChIP-qPCR analysis depicted HDAC3 promoter enrichment for the MMP3/13 gene in IPMK F/F, IPMK^Δ^*^IEC^* and IPMK^Δ^*^IEC^* mice pretreated with oral InsP_6_. Data represents three experimental replicates (n=3).

**Extended Figure 5II:** (A) IPMK KO cell lysate and also conditioned media show MMP 13 and 3 enrichment. Densitometric analysis of western blot from Fig 5 G for (B) MMP3; (C) MMP13; (D) ZO-1, Occludin (E), and (F) Claudin 2. Pathway analysis using the Metascape tool for RNA-seq data shows (G) upregulated genes in IPMK^Δ^*^IEC^* compared to IPMK^F/F^ and (H) downregulated pathways. (I) Relevant enriched gene set from GSEA of RNA-seq datasets for *IPMK^F/F^ vs IPMK*^Δ*IEC*^ of the mouse model, using the MSigDB C2:CP:Reactome subcollection.

**Extended Figure 6: (A)** Western blot analysis of colonic crypts diminished IPMK expression after 24 hours of 3% DSS treatment, which was not restored by oral InsP6 pretreatment. While IPPK expression remained unchanged in both treated and untreated groups. Lamin served as the loading control. Data are representative of three independent experiments (n = 4). **(B)** ChIP-qPCR analysis depicted HDAC3 promoter enrichment for MMP3/13 gene in IPMK *^F/F^,* IPMK^Δ^*^IEC^* and IPMK^Δ^*^IEC^* mice pretreated with oral InsP_6_. Data represents three experimental replicates (n=3).

**Extended Figure 7.** (A) Average body weight of wild-type mice. (B) H&E staining of the colon from untreated, DSS, and DSS+InsP6-treated mice. n=3.Scale bar represents 60 μm (C) Body weight plot of IPMK^F/F^ and IPMK^ΔIEC^ mice. (D) Colon length of IPMKF/F and *IPMK*^ΔIEC^ mice after DSS treatment. n=3. (E) H&E staining of the colon from IPMK^F/F^ and IPMK^ΔIEC^ mice after DSS and DSS+InsP6 treatment. n=3. Scale bar represents 60 μm (F) MiniMUGA mouse backcross genotyping analysis by Transnetyx was performed using two *IPMK^F/F^* mice, confirming that the mice were backcrossed for 10 generations to the C57BL/6 strain, achieving a congenic background of 97% or higher.(n=2)

