## Extended File for "Phytic Acid (InsP_6_) Activates HDAC3 Epigenetic Axis to Maintain Intestinal Barrier Function"

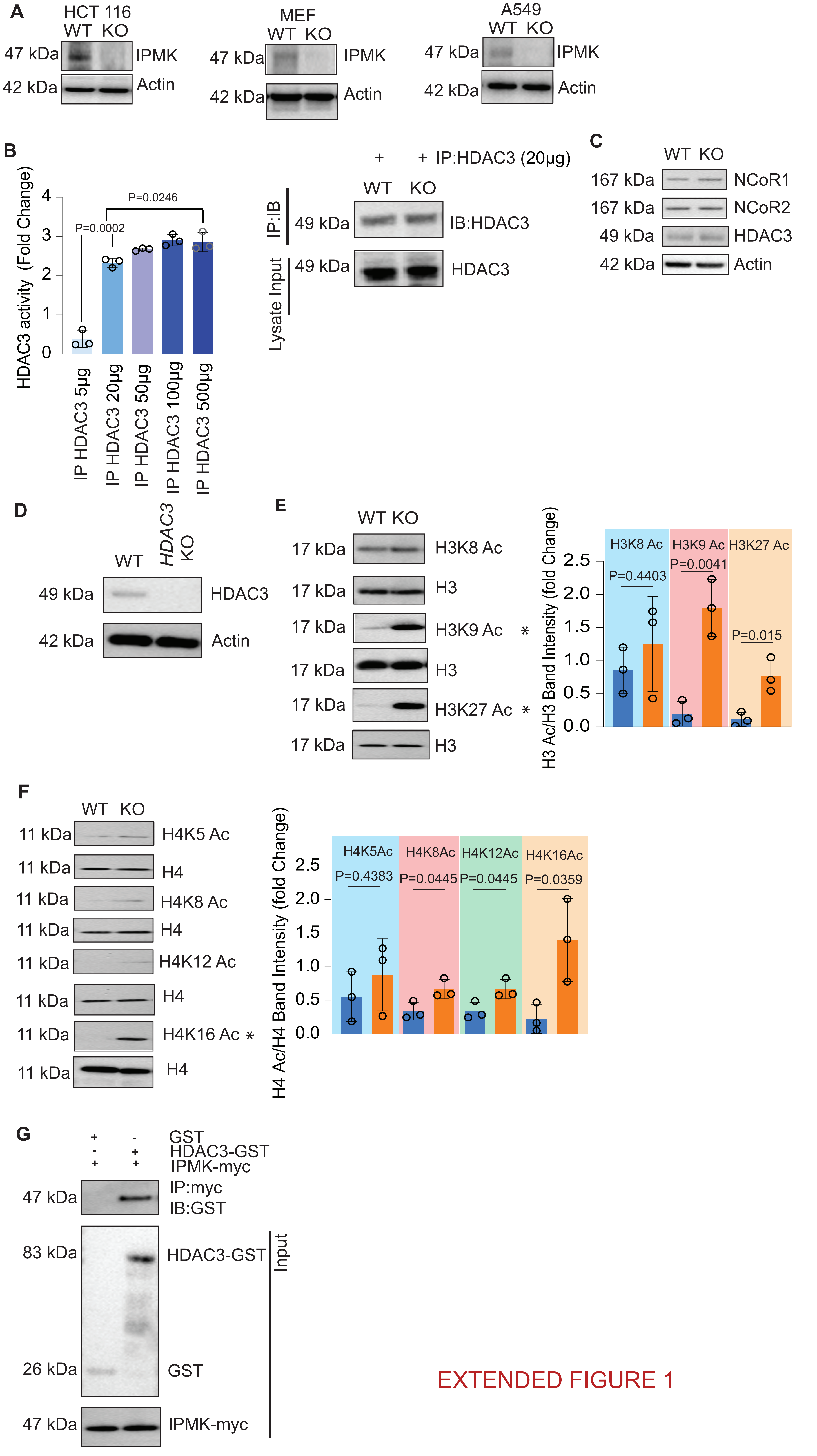


EXTENDED FIGURE 1

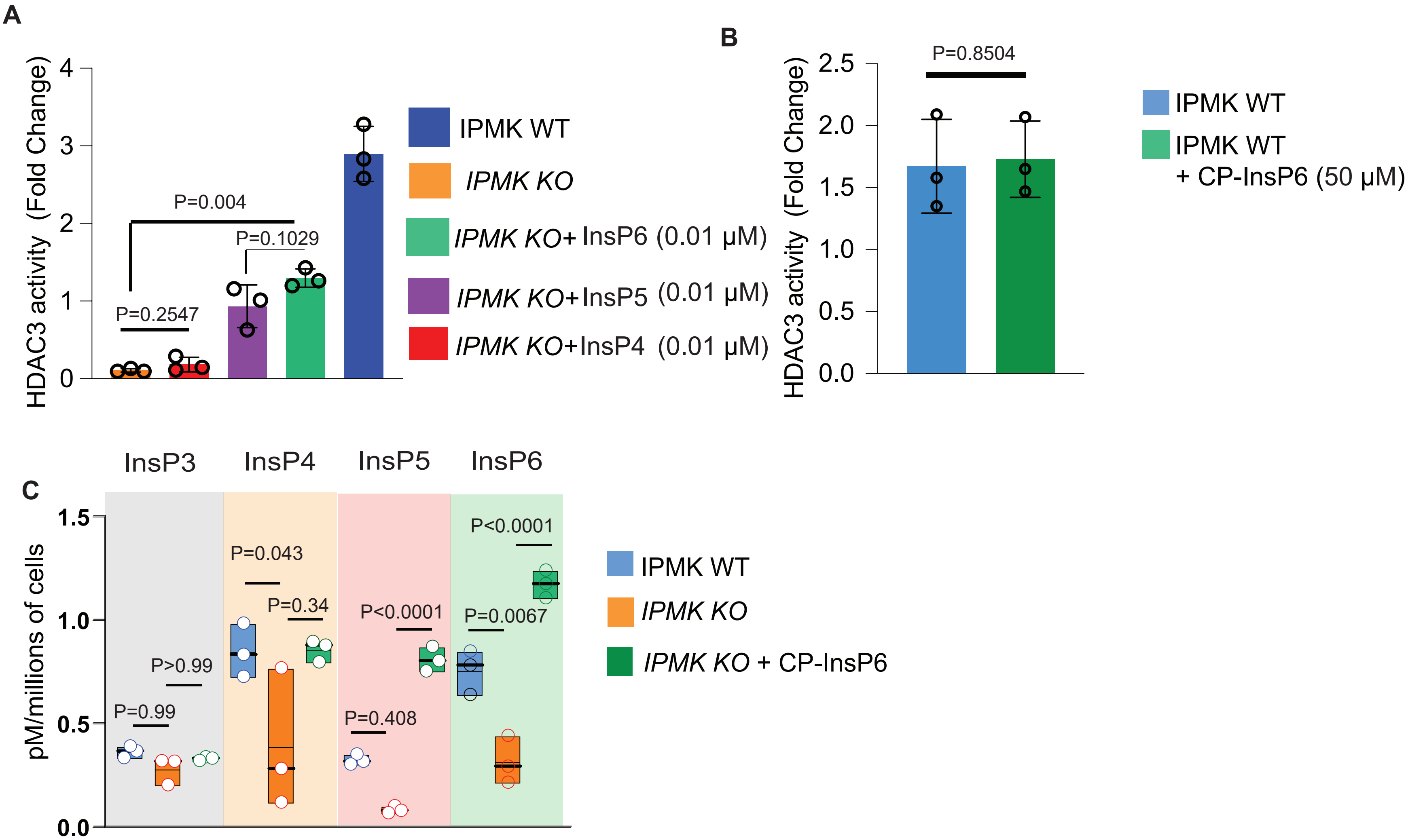


### EXTENDED FIGURE 2

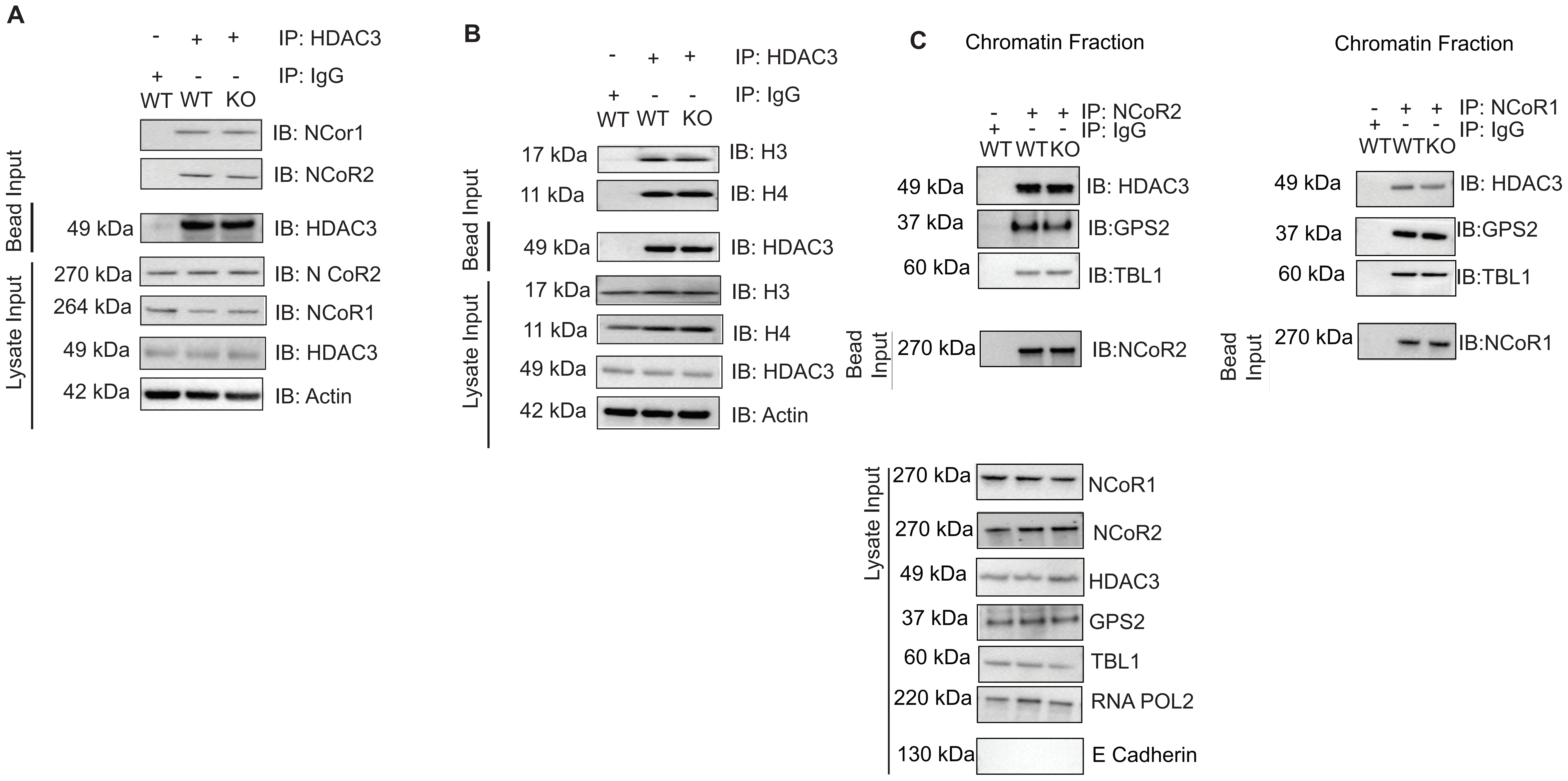


Extended Figure 3 I

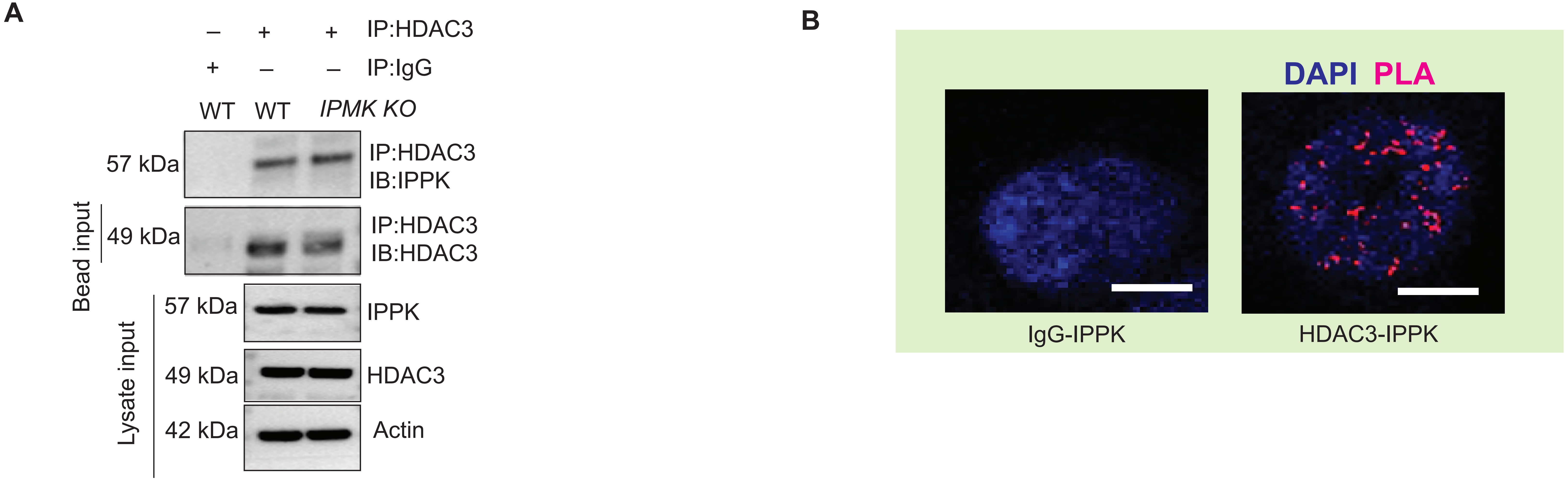


Extended Figure 3 II

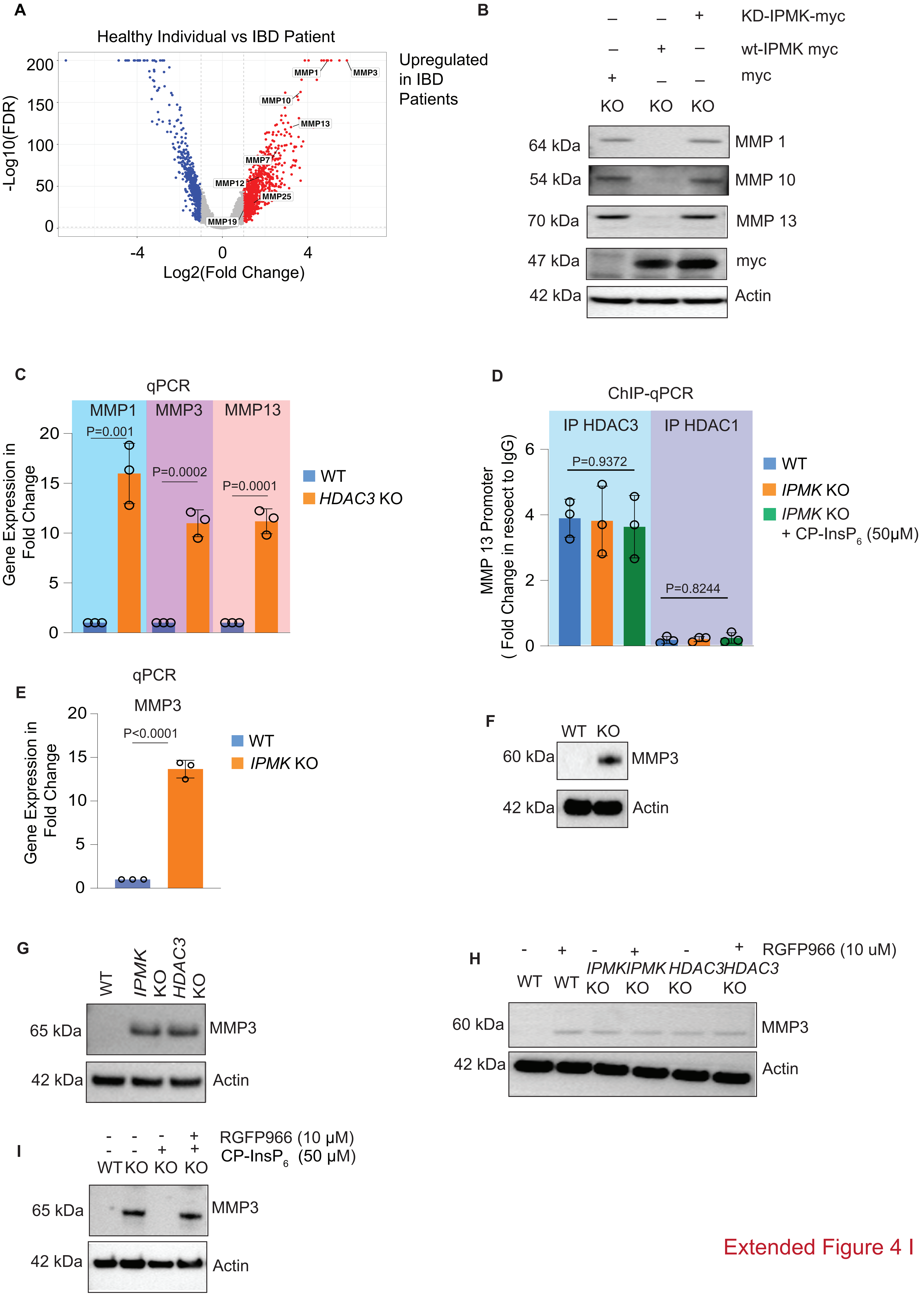


Extended Figure 4 I

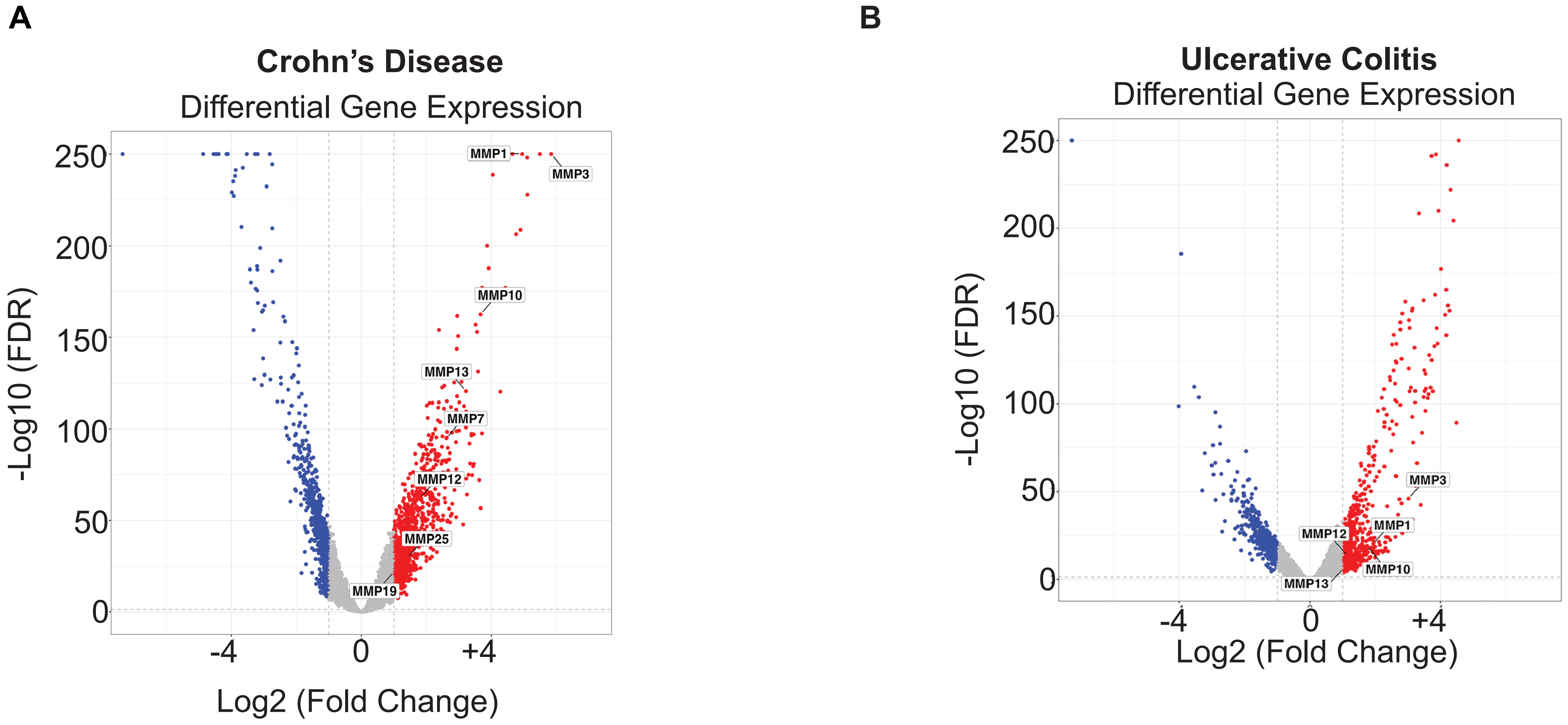


Extended Figure 4 II

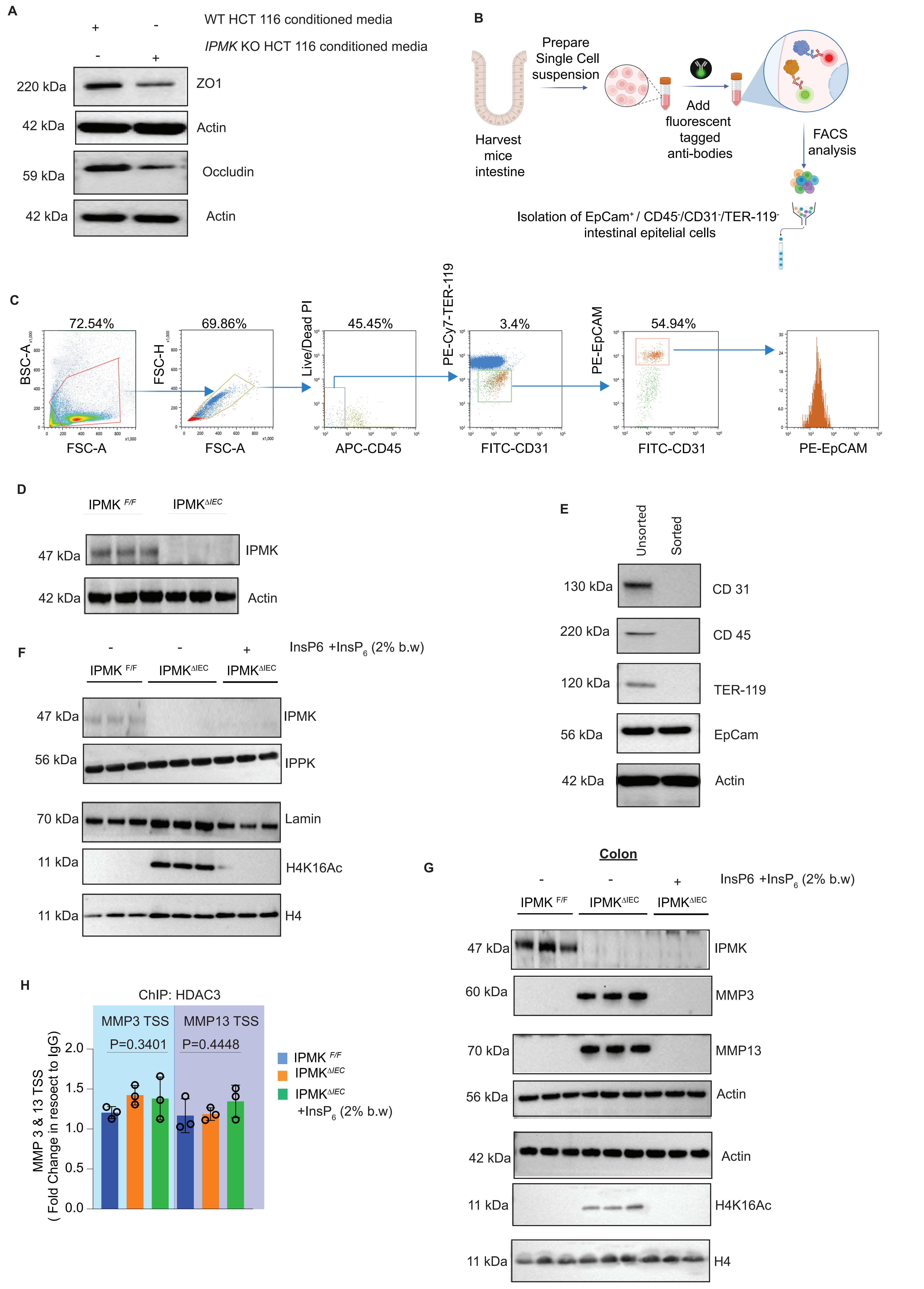


### EXTENDED FIGURE 5 I

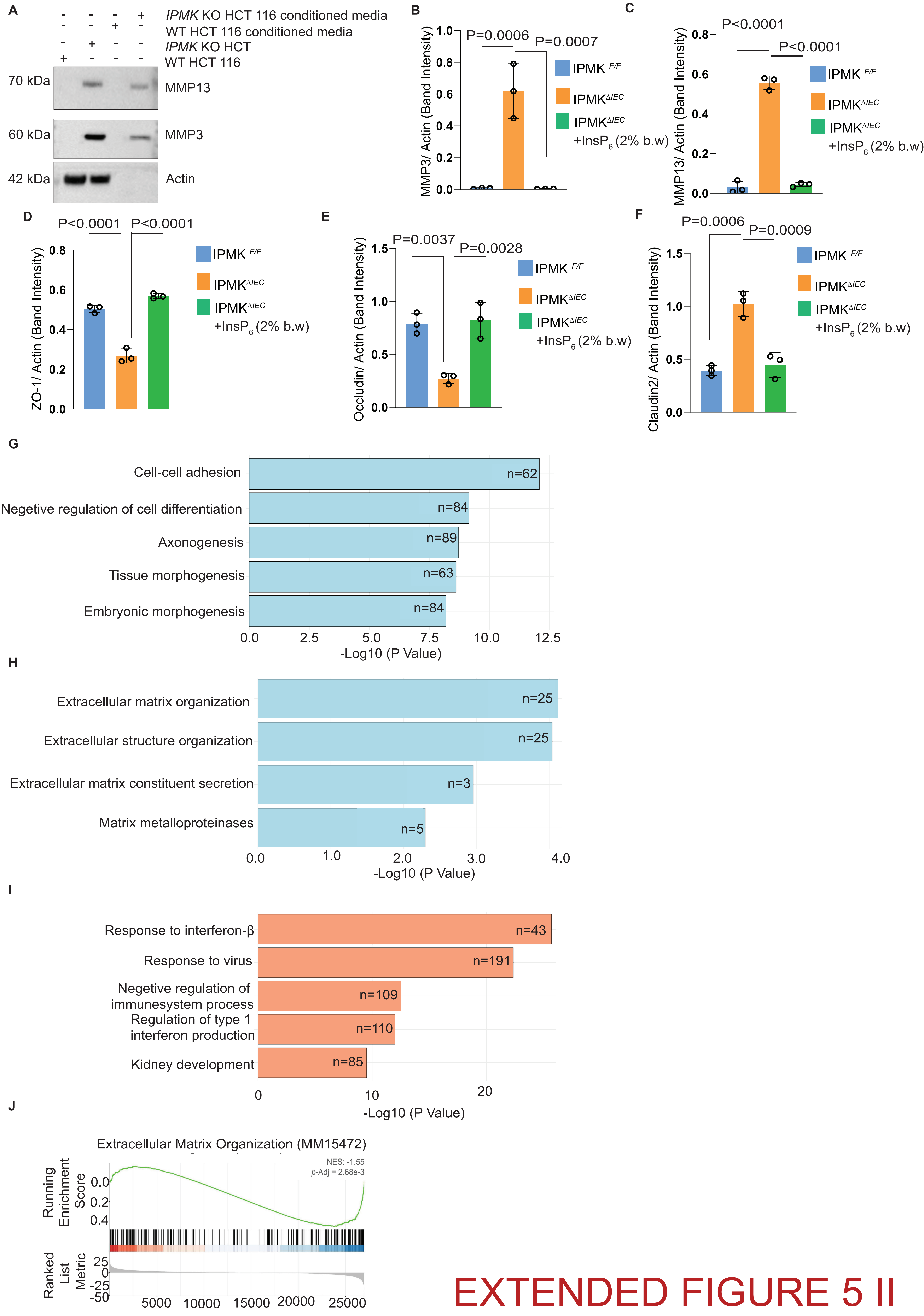


EXTENDED FIGURE 5 II

A

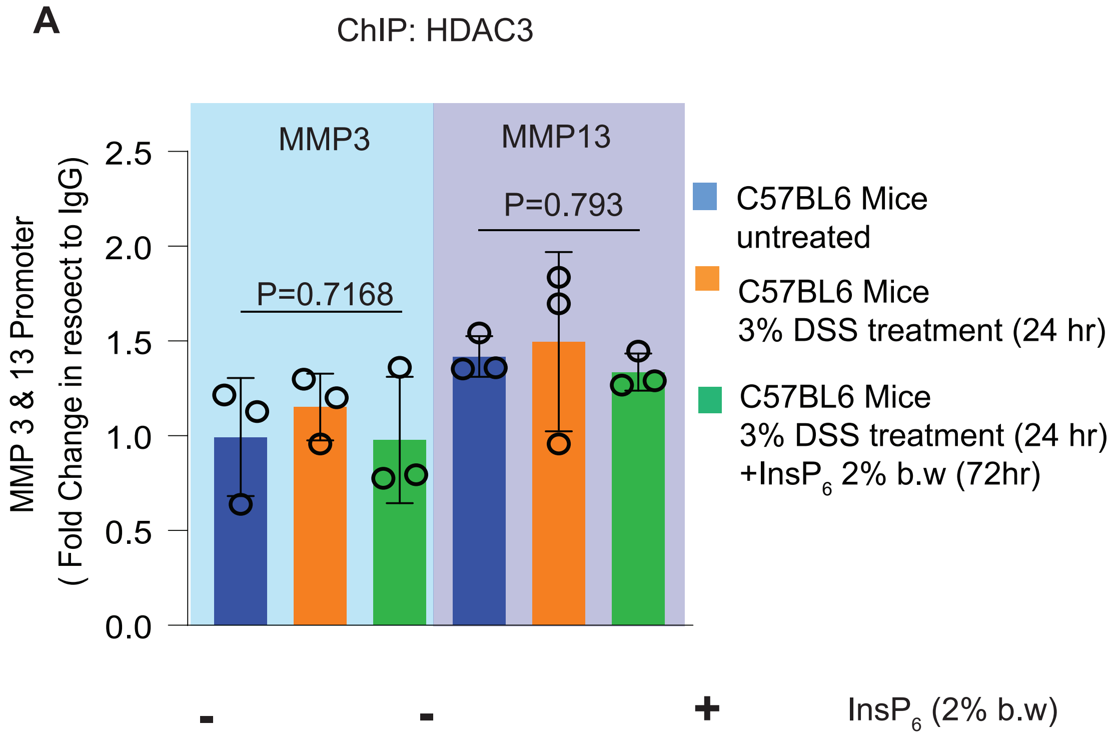


B

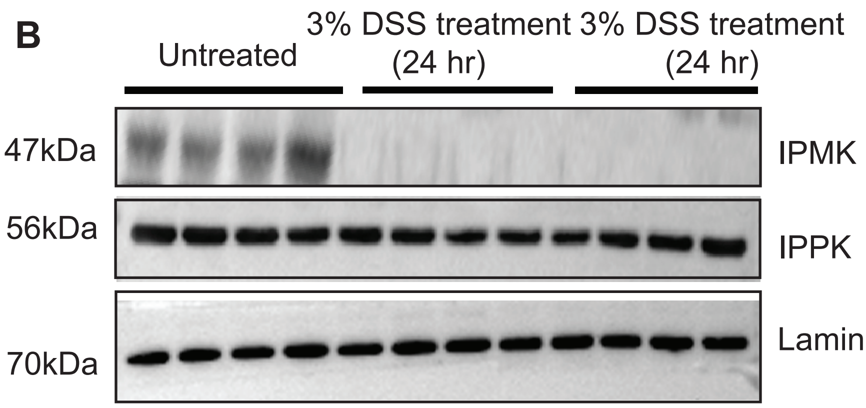


### Extended Figure 6

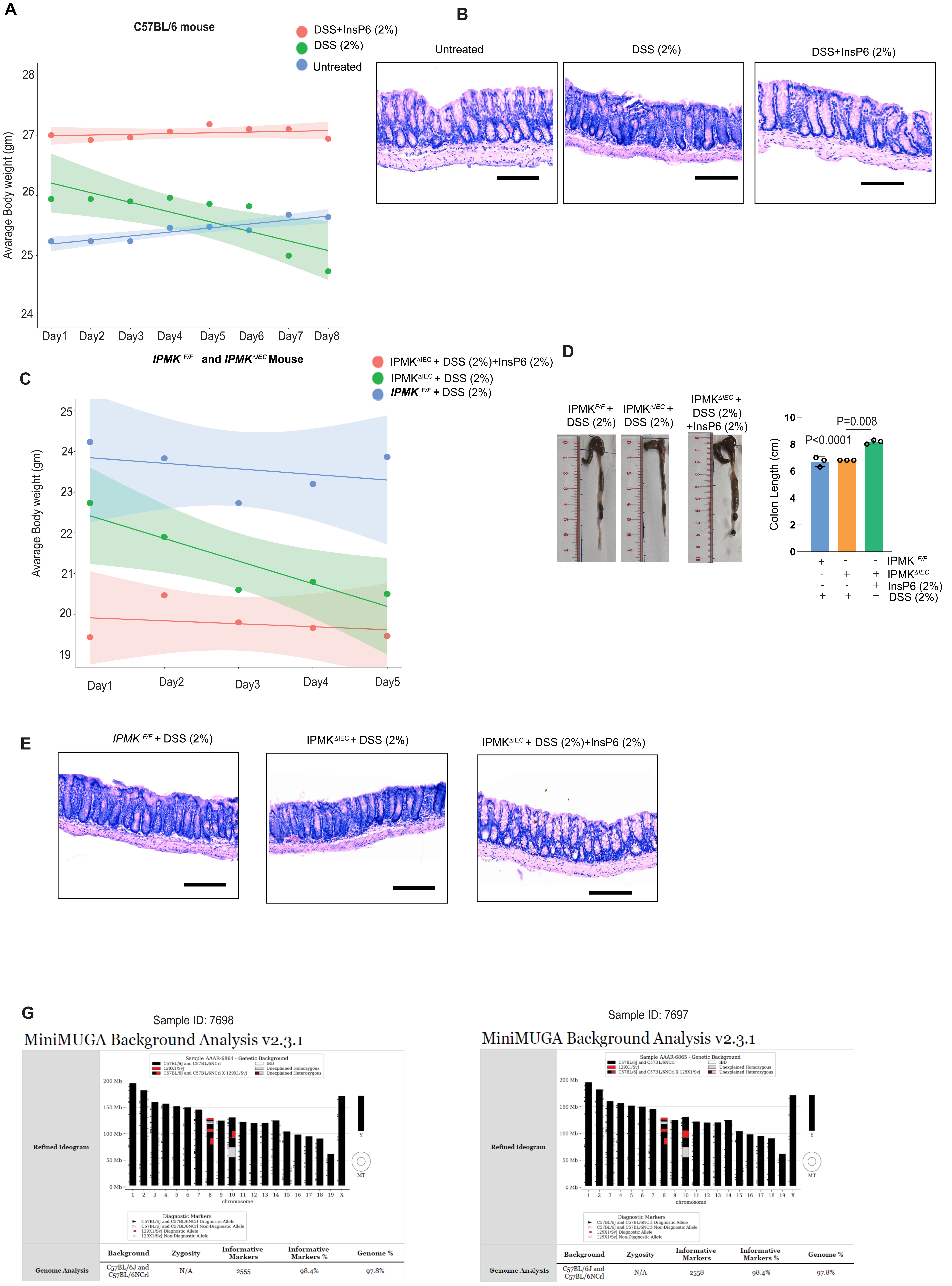


Extended Figure 7
